# Accuracy of experimentally estimated muscle properties: Evaluation and improvement using a newly developed toolbox

**DOI:** 10.1101/2025.09.29.678508

**Authors:** Edwin D.H.M. Reuvers, Dinant A. Kistemaker

## Abstract

The mechanical behaviour of a muscle-tendon complex depends on properties such as the force-length relationships, the force-velocity relationship, and the excitation dynamics. Quick-release and step-ramp experiments are commonly used to estimate these properties. The accuracy of these methods is unclear, as the actual values of these properties are unknown in experiments on real muscle. We conducted a modelling study using a Hill-type muscle-tendon complex model with literature-derived parameter values and simulated quick-release, step-ramp, and isometric experiments. From the simulated experiments, we assessed how accurately the model’s parameter values could be retrieved. Using a method traditionally used in literature, the series elastic element stiffness was underestimated by ~35%, due to the incorrect assumption that muscle fibres do not shorten during quick releases. Consequently, this yielded an overestimation of the excitation dynamics activation time constants of ~20%. We developed an improved method that accounted for muscle fibre length shortening during quick releases. Using our improved method, all parameter values closely matched their actual values. A sensitivity analysis showed that the most critical parameters were robust to perturbations in experimental data. Lastly, we compared Hill-type MTC model predictions against *in situ* data from three rat m. gastrocnemius medialis muscles. Predictions based on parameters from the improved method showed closer agreement than those based on the traditional method — both for quick-release, step-ramp, and isometric experiments, as well as for independent stretch-shortening cycles. In conclusion, the improved method enables more accurate estimates of muscle-tendon complex properties, addressing limitations of the traditionally used method.

## 1 Introduction

In the early 20th century, A.V. Hill laid the foundation of muscle mechanics by recognising that muscles contain not only muscle fibres (the contractile element; CE) but also connective tissue, both parallel to the muscle fibre (the parallel elastic element; PEE) and in series with the muscle fibre (the tendon or serial elastic element; SEE). Together these three elements form the muscle-tendon-complex (MTC). The contraction dynamics describe how CE force emerges from the interplay between the force-length properties of CE, PEE and SEE, and the force-velocity properties of CE. In addition, CE force depends on the active state (the relative amount of *Ca*^2+^ bound to troponin C, Ebashi and Endo, 1968). Active state, in turn, depends on activation via the excitation dynamics. Consequently, the mechanical behaviour of MTC arises from the intricate interplay between the contraction and the excitation dynamics.

In his famous (1938) paper A.V. Hill performed so-called’isotonic quick-release experiments’ using the now-renowned Levin-Wyman lever (Levin and Wyman, 1927) to estimate the CE force-velocity relationship. In these experiments, MTC was attached to a lever and maximally stimulated. When SEE force plateaued (i.e., MTC was isometrically delivering force), a magnet was released from the lever causing a sudden drop in the force acting on the MTC. As SEE force remained constant right after this drop in SEE force, SEE length remained constant and therefore all MTC length changes were attributed to CE length changes. The CE velocity corresponding to the constant SEE force after the drop was then calculated as the maximum rate of change of MTC length over time. Consequently, each isotonic quick-release experiment contributed a single data point of the CE force-velocity relationship. Nowadays, servomotors are typically used in experiments on isolated MTCs to control MTC length (changes). In contrast to the force-controlled experiments with the Levin-Wyman lever, servomotor control the MTC length over time (see Figure S1). In order to estimate the CE force-velocity relationship using a servomotor, Cecchi et al. (1978) introduced’step-ramp experiments’. In these experiments, MTC is kept isometrically under maximal stimulation until SEE force plateaus. Then, a rapid shortening of MTC length is imposed (the’step’), immediately followed by a constant-velocity shortening of the MTC (the ‘ramp’). This experimental protocol results in a brief plateau in SEE force just after the ramp. During this brief period, SEE length remains constant and therefore CE velocity equals MTC velocity. Thus, step-ramp experiments with servomotors serve as a valid alternative for isotonic quick-release experiments to estimate the CE force-velocity relationship.

In addition to estimating the CE force-velocity relationship using isotonic quick-release experiments, the research group of A.V. Hill also used these experiments to approximate SEE stiffness. The underlying idea was that the drop in MTC force was so fast that, due to contraction dynamics, CE had no time to shorten and all MTC shortening could thus be directly attributed to SEE shortening. Consequently, this experiment allowed to directly relate SEE length changes to changes in SEE force and therefore to estimate SEE stiffness. As mentioned above, in isotonic quick-release experiments, a sudden drop in force acting on the MTC causes a rapid change in MTC length. While in these lever-based experiments MTC force is controlled, in experiments using a servomotor MTC length is controlled. To estimate SEE stiffness with a servomotor setup, the opposite approach is taken: a rapid change in MTC length is imposed by the motor, resulting in a rapid change in SEE force. Somewhat confusingly, these servomotor-based experiments are also referred to as ‘quick-release experiments’, even though their approach differs fundamentally from that of an isotonic quick-release experiment. In the remainder of this paper, the term quick-release experiments will exclusively refer to servomotor-based length-controlled quick-release experiments. In quick-release experiments using a servomotor, the drop in MTC length takes several milliseconds (e.g., approximately 10 ms with the commonly used Aurora Scientific’s 300C series motor). As CE shortens within this short time period, the assumption that MTC length changes are exclusively due to SEE length changes may not be entirely valid. This raises the question to what extent SEE stiffness can be accurately estimated based on quick-release experiments using servomotors.

A.V. Hill already acknowledged in (1950) that CE shortens during the brief period in which SEE force decreases – even for his isotonic quick-release experiments using the Levin-Wyman lever – and introduced a method to correct for this CE shortening. In this method, the CE shortening that occurred due to the quick-release was estimated based on the CE force-velocity relationship of the MTC investigated. Thus, additional experimental work – such as performing step-ramp experiments and estimating the CE force-velocity relationship – are required to correct for CE shortening during the quick-release. Zandwijk et al. (1997) and Lemaire et al. (2016) developed similar methods to correct for CE shortening. As expected, Zandwijk et al. (1997) showed that this method results in higher estimates of SEE stiffness than with the method without correcting for CE shortening. Nevertheless, the accuracy of SEE stiffness estimates remains uncertain, as the actual SEE stiffness is unknown in experiments, making it infeasible to compare the estimated value against the actual one. Consequently, it remains unclear whether the additional complexity and experimental work required for the improved method are truly justified in comparison with the simpler traditional method, especially when the only interest is SEE stiffness.

In experimental studies in which MTC properties are estimated, SEE stiffness is typically estimated first in order to discriminate between SEE stiffness and the other properties, such as those of the CE force-length relationship. In experiments, CE length is unknown and therefore the properties of the CE force-length relationship cannot be directly estimated based on experimental data. Typically, the CE force-length properties are estimated based on the measured MTC force-length relationship (e.g., Blümel et al., 2012; Lemaire et al., 2016). However, different combinations of CE and SEE properties can yield almost identical MTC force-length relationships. Consequently, inaccuracies in SEE stiffness estimates can propagate to errors in the estimation of the CE force-length properties. While predictions under isometric conditions may remain reasonably accurate, these errors can lead to substantially different mechanical behaviour during dynamic contractions such as stretch-shortening cycles (see Figure 1). As such, incorrect estimation of SEE stiffness might (partially) explain difference between predictions derived by a Hill-type MTC model and experimental results. In sum, the question arises to which extent the estimates of other MTC properties are affected by inaccurate SEE stiffness estimates and whether more accurate SEE stiffness estimates improves predictions derived by a Hill-type MTC model.

**Figure 1:**
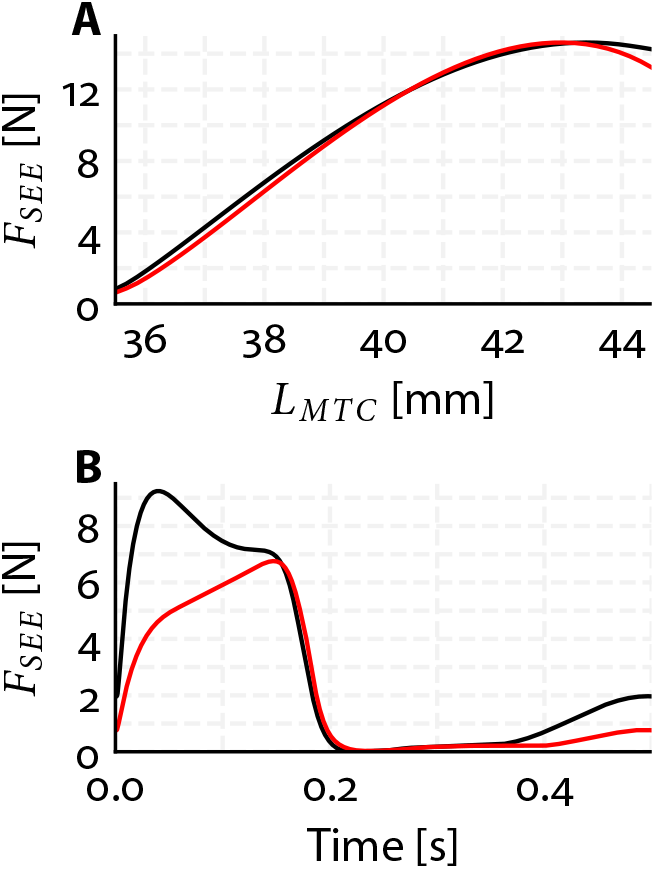
Effect of a 2.5-fold difference in SEE stiffness on simulated mechanical behaviour. This example shows that some contractions may lead to similar mechanical behaviour across distinct parameter sets, others may differ substantially.

The potential effect of error propagation also extends to the estimation of the excitation dynamics properties. In the absence of histochemical data, excitation dynamics properties are commonly estimated by fitting predicted SEE force of the Hill-type MTC model to experimental data (e.g., Blümel et al., 2012; Lemaire et al., 2016). As SEE force over time obviously depend on the contraction dynamics properties (e.g., the force-length relationships and the force-velocity relationship), the estimation of the excitation dynamics properties is not independent of the estimation of the contraction dynamics properties. Consequently, inaccuracies in SEE stiffness estimates can propagate to inaccuracies in estimated excitation dynamics properties. This interdependence raises an additional question of what experimental data should be used in order to minimise this potential error propagation.

The primary objective of this study was to systematically evaluate the accuracy of estimating contraction and excitation dynamics properties on the basis of data collected in commonly used experiments. Specifically, we aimed to compare the results of a method that does not correct for CE shortening in quick-release experiments (the ‘traditional method’) with a method that does correct for CE shortening (the ‘improved method’). As explained earlier, the limitation of relying exclusively on an experimental approach is that the actual properties are unknown, which renders a reliable assessment of the accuracy of the estimated properties not feasible. To circumvent this problem, we conducted a modelling study using a Hill-type MTC model. First, we obtained three different sets of parameter values of a Hill-type MTC model from existing literature. Using these parameter values, we simulated data of quick-release, step-ramp and isometric protocols mimicking experiments on isolated MTCs using servomotors. We then investigated how accurately we could retrieve the model’s parameter values (with respect to their ‘actual’ value). In addition, we employed a comprehensive sensitivity analysis to assess the sensitivity of parameter values against perturbations in experimental data and to examine the interdependency of the estimated properties. The secondary objective of this study was to evaluate whether predictions from a Hill-type MTC model using parameter values estimated with the improved method better matched experimental data than predictions using parameter values estimated with the traditional method. To this end, we used *in situ* data from experiments on three rat m. gastrocnemius medialis. We estimated the contraction and excitation dynamics properties using both the traditional and the improved method, and compared the resulting model predictions — based on each parameter set — to experimental force data. Overall, this study provides insights into the accuracy of existing commonly used methods for contraction and excitation dynamics property estimation and their influence on predictions of mechanical behaviour derived by a Hill-type MTC model. The approach that we designed in this study for estimating contraction and excitation dynamics properties based on data of quick-release, step-ramp and isometric experiments using servomotors, is made available as an open-source toolbox (https://github.com/edwinreuvers/mp-estimator).

## 2 Methods

The primary objective of this study was to assess the accuracy of estimating contraction and excitation dynamics properties. For this purpose, we employed a Hill-type MTC model to simulate MTC mechanical behaviour, as detailed in Section 2.1. Using parameter values obtained from existing literature, we simulated data of quick-release, step-ramp, and isometric experiments, mimicking experiments on isolated MTCs with servomotors, as described in Section 2.2.1. We then estimated the contraction and excitation dynamics parameter values using two methods: one with (the ‘traditional method’) and one without (the ‘improved method’) a correction for CE shortening due to the quick-release (see Section 2.3). Additionally, we conducted comprehensive sensitivity analyses to assess the accuracy of the improved method, as outlined in Section 2.4. The secondary objective was to evaluate predictions derived by a Hill-type MTC model using two sets of parameter values: one estimated by the traditional method, the other by the improved method. For this purpose, we used *in situ* data (see Section 2.2.2) from three rat m. gastrocnemius medialis. Predictions were assessed against quick-release, step-ramp, and isometric experiments, as well as independently measured stretch-shortening cycles. This allowed us to investigate whether the improved method enhanced the predictions derived by a Hill-type MTC model.

### 2.1 Muscle model

#### 2.1.1 Contraction dynamics

We employed a Hill-type MTC model consisting of a contractile element (CE) and a parallel elastic element (PEE), which were both in series with a serial elastic element (SEE), as depicted in Figure 2. CE represented the contractile element of the muscle fibres, while PEE and SEE represented the tissues arranged in parallel and in series with the muscle fibres, respectively. Given the negligible small mass of MTC compared to the forces typically delivered by CE, PEE, and SEE, we simplified the model by neglecting the second-order dynamics of the system:

**Figure 2:**
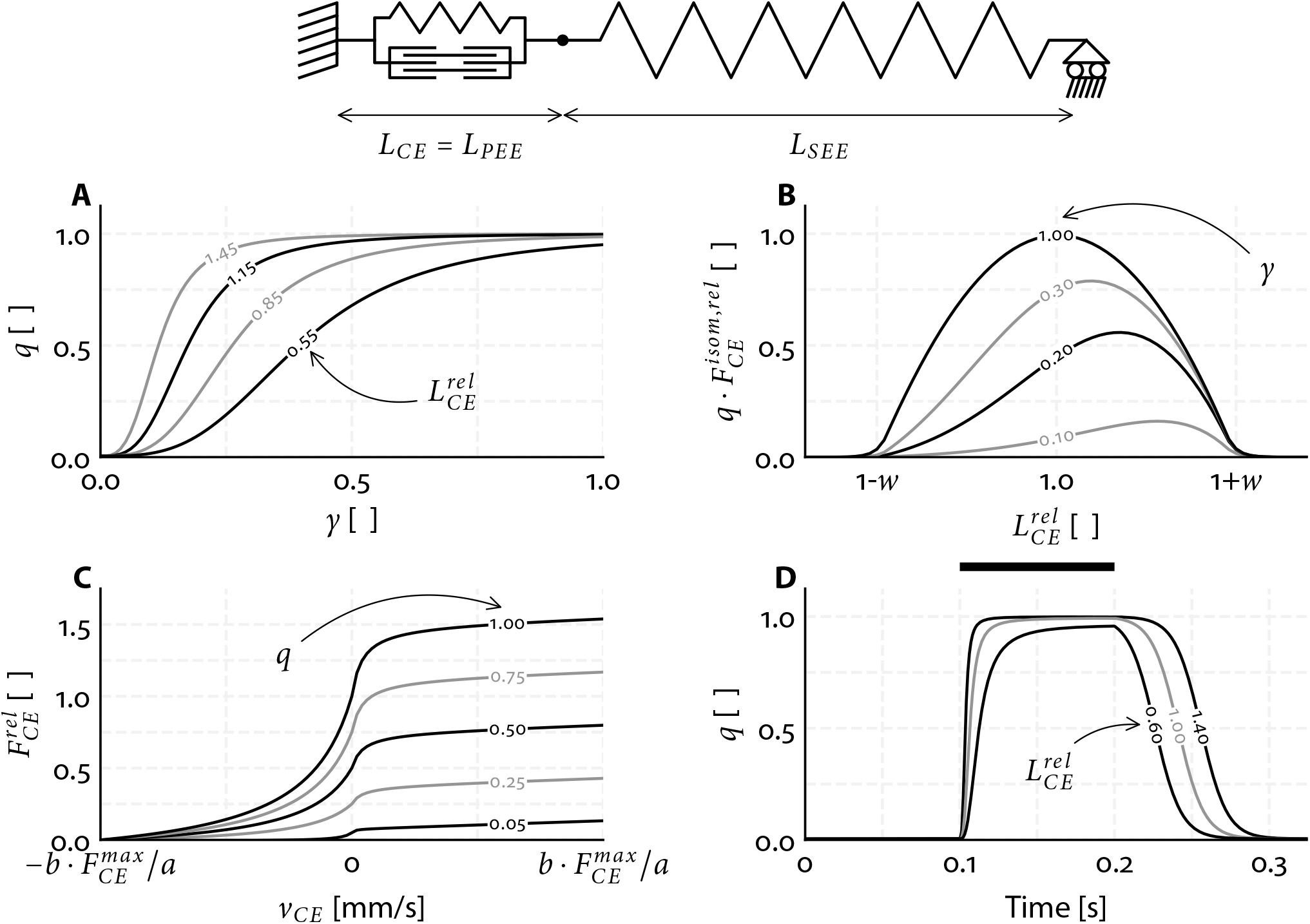
Aspects of the Hill-type MTC model used, illustrated at the top. *L*_*CE*_, *L*_*PEE*_ and *L*_*SEE*_ denote the CE, parallel elastic element (SEE) and serial elastic element (SEE) length. CE represents the contractile part of the muscle fibres, while PEE and SEE represent all elastic tissue in parallel or in series, respectively, with CE. In the Hill-type MTC model, CE force depends on active state (A), CE length (B) and CE velocity (C). The effect of CE stimulation on active state (*q*) is illustrated in D. A) The relationship between normalised free *Ca*^2+^ concentration between the myofilaments (*γ*) and *q. q* also depends on relative CE length 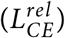. B) The product of *q* and the normalised active CE force-length relationship for different values of *γ*. C) The CE force-velocity relationship for different values of *q*. D) *q* over time before, during and after CE stimulation for 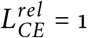. CE stimulation is maximal during the period indicated by the black bar and ‘off’ elsewhere.

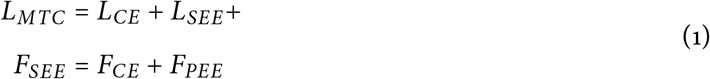

*L*_*MTC*_, *L*_*CE*_, *L*_*SEE*_ and *L*_*PEE*_ denote MTC, CE, PEE and SEE length respectively, while *F*_*SEE*_, *F*_*CE*_, and *F*_*PEE*_ denote the CE, PEE and SEE force. PEE and SEE were assumed be purely elastic and were modelled as quadratic springs (Zajac, 1989):

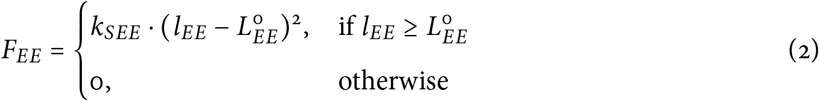

*L*_*EE*_ denotes the length of either PEE or SEE and 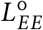 the slack length of either PEE or SEE. *k*_*EE*_ denotes a parameter that scales the stiffness of either PEE or SEE.

Isometric CE force depends on CE length (Blix, 1892; Blix, 1893; Hill, 1925). We simplified the classic sarcomere force-length relationship (Gordon et al., 1966; Walker and Schrodt, 1974) to a second order polynomial, which describes the force-length relationship reasonably well (Bobbert et al., 1990; Woittiez et al., 1984):

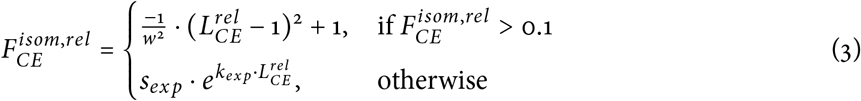

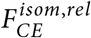 denotes the CE isometric force normalised by 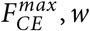 determines the width of the CE force-length relationship (for 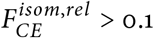) and 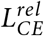 denotes CE length normalised by CE optimum length (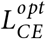, the CE length at 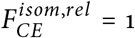). Exponential tails were added to the isometric CE force-length relationship such that it had a continuous first derivative with respect to 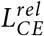 (see Figure 2). The parameter values of *s*_*exp*_ and *k*_*exp*_ were determined separately for the ascending and descending limb of the CE force-length relationship.

The concentric CE force-velocity was modelled according to Hill (1938). The original description was adjusted as suggested by Soest and Bobbert (1993) to account for situations where the active state (*q*, the relative amount of *Ca*^2+^ bound to troponin C, Ebashi and Endo, 1968) is not maximal:

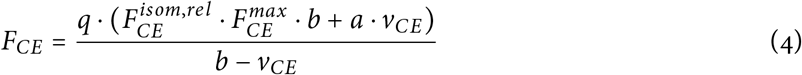

*F*_*CE*_ denotes the CE force and *v*_*CE*_ denotes the CE velocity. *a* is a parameter that defines the curvature of the concentric CE force-velocity relationship and defines together with *b* and 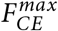 the maximum shortening velocity 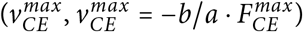. The maximum shortening velocity is scaled by 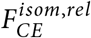 due to the formulation of the CE force-velocity relationship (see Equation 4). However, the maximum shortening velocity has been reported to be more or less constant above optimum CE length (Gordon et al., 1966; Stern, 1974). Based on this observation, the parameter *a* was scaled by 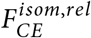 above optimum CE length 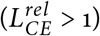 to make the maximum shortening velocity independent of CE length above optimum CE length. The parameter *b* was scaled with *b*_*scale*_ for low levels of active state in order to make the maximal contraction velocity dependent on the active state (Petrofsky and Phillips, 1981). The original description of this scaling by Soest and Bobbert (1993) was somewhat reformulated, such that *b*_*scale*_ was continuously differentiable with respect to *q*:

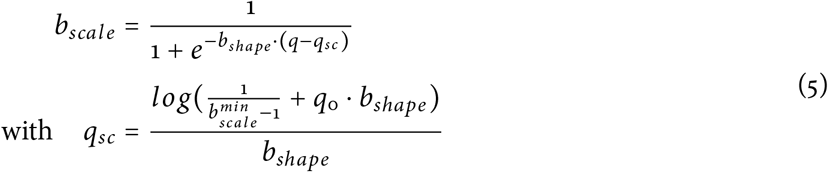

*q*_0_ denotes the minimum value of 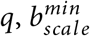 denotes the minimal scale factor of *b* (i.e., the minimal value of *b*_*scale*_) and *b*_*shape*_ defines the steepness of the relation between *b*_*scale*_ and *q*. The value of *b*_*shape*_ was obtained by minimising the least square error between the formulation of Soest and Bobbert (1993) and the reformulated version of the relation between *b*_*scale*_ and *q*.

The eccentric part of the CE force-velocity relationship was modelled as a slanted hyperbola function (Soest et al., 2005) of the following form:

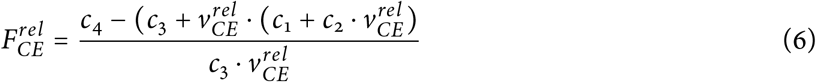

*c*_1_, *c*_2_, *c*_3_ and *c*_4_ denote parameters that define the shape of the slanted hyperbola and depend on 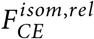 and *q*. These parameters were chosen such that (1) the concentric and eccentric curves were continuous in the point 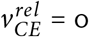; (2) the ratio between the eccentric and concentric derivatives of 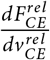 in the point 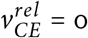 was defined by *r*_*slope*_; (3) the oblique asymptote of the eccentric curve was defined by *F*_*asymp*_ and (4) the slope of the slanted asymptote was defined by *r*_*as*_.

#### 2.1.2 Excitation dynamics

The excitation dynamics were modelled in two steps. The first step, to which we refer as the calcium dynamics, related the rate of change in normalised free *Ca*^2+^ concentration between the myofilaments (*γ*) to normalised *CE* stimulation (*STIM*) and the normalised free *Ca*^2+^concentration between the myofilaments itself (Hatze, 1981, pp 31-42):

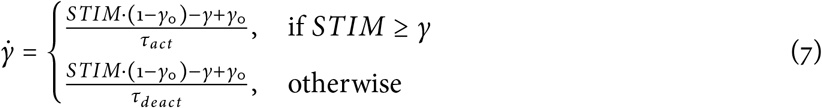

*γ*_0_ denotes the minimum value of *γ* and had a arbitrary small value such that Equation 8 was always solvable. *τ*_*act*_ and *τ*_*deact*_ are both time constants of the first-order differential equations describing the calcium dynamics. The second step involved the relation between active state (*q*) and *γ*, to which we refer as the *q* − [*Ca*^2+^] relation. It is well-known that muscle fibres become increasingly sensitive to [*Ca*^2+^] as their length increases (Kistemaker et al., 2005; Rack and Westbury, 1969; Stephenson and Williams, 1982). Consequently, *q* was also made dependent on 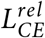. The *q* − [*Ca*^2+^] relation was modelled according to Hatze (1981, pp 31-42), but was mathematically reformulated to ensure that the parameter values were physiologically meaningful and to facilitate its application for Optimal Control (e.g., Kistemaker et al., 2023):

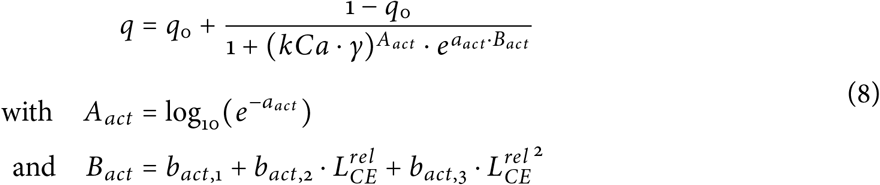

*kCa* relates *γ* to the actual *Ca*^2+^ concentration between the myofilaments and *B*_*act*_ denotes the *pCa*^2+^ level at which *q* = 0.5. *B*_*act*_ depended on 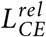 (Equation 8), while parameter *a*_*act*_ determined the steepness of the relation between *γ* and *q*.

### 2.2 Simulated and *in situ* data

#### 2.2.1 Simulated data

To simulate quick-release, step-ramp, and isometric experiments, we used three parameter sets of a Hill-type MTC model from existing literature for three rat m. gastrocnemius medialis (GM1, GM2, and GM3; see Table S1).

##### Quick-release experiments

Each quick-release experiment consisted of an isometric phase until SEE force plateaued, followed by a rapid (step) change (10 ms) in MTC length and then followed by another isometric phase (see Figure 3A). CE stimulation was maximal (i.e., *STIM* = 1) during the first isometric phase and continued at this maximal level until it was switched off shortly after the step change in MTC length occurred. The step change in MTC length was 0.2 mm and was chosen such that the resulting change in SEE force was about 5-10% of 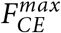. To prevent irreversible damage to muscle fibres, the maximal MTC length used in experiments is typically not far above the MTC length that yields maximal isometric SEE force 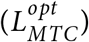. Accordingly, we simulated quick-release experiments at various initial MTC lengths, ranging from a very short MTC length (i.e., a length yielding very low isometric SEE force) and at every 1 mm increment up to a MTC length that was maximal 3 mm above 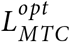.

**Figure 3:**
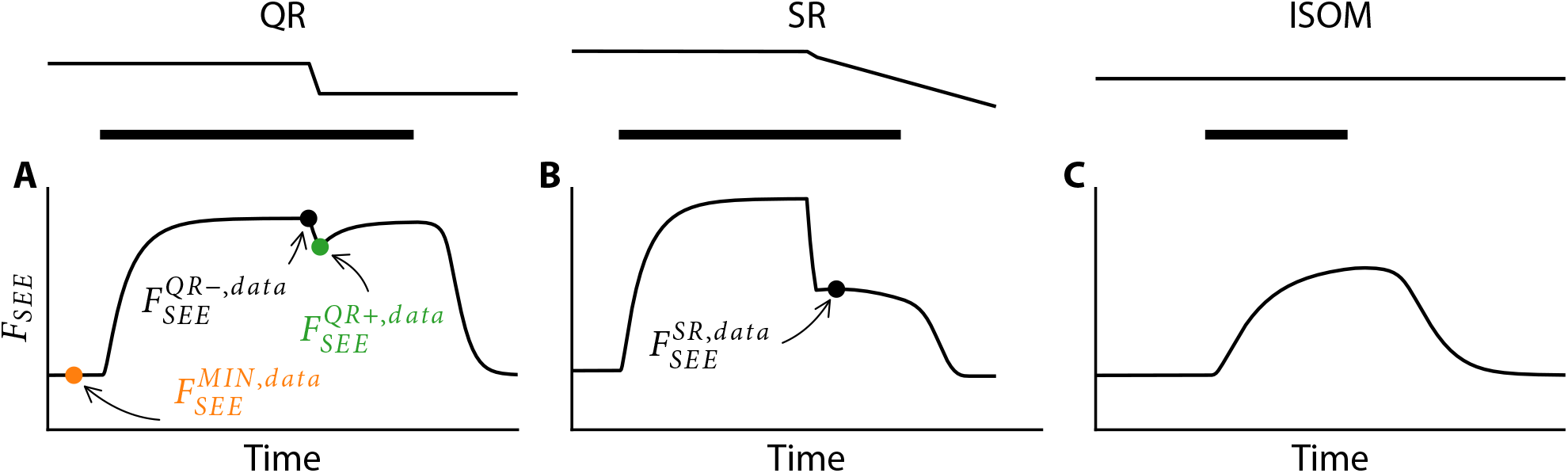
Example of simulated data of quick-release (A), step-ramp (B) and isometric (C) experiments. Top: MTC length over time. Bottom: SEE force over time. CE stimulation is maximal during the period indicated by the black bar, and ‘off’ elsewhere. For each experiment, we obtained specific datapoints of SEE force and the corresponding MTC length, which were used to estimate contraction and excitation dynamics parameter values.

##### Step-ramp experiments

Each step-ramp experiment consisted of an isometric phase slightly above (± 0.5 mm) 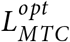, followed by a rapid (step) change (10 ms) in MTC length and then a constant MTC velocity ramp (see Figure 3B). CE stimulation was maximal (i.e., *STIM* = 1) during the first isometric phase and continued at this maximal level until it was switched off 0.1 s after the step change in MTC length occurred. The change in MTC length of this step and the constant MTC velocity of the ramp were chosen such that this resulted in a more or less a constant SEE force (and therefore constant CE force) at the beginning of the ramp. We simulated 9 step-ramp experiments with different combinations of step sizes and constant MTC velocity ramps, to cover a substantial part of the concentric CE force-velocity relationship.

##### Isometric experiments

Isometric experiments were simulated for a combination of different MTC lengths (about 0, −2, −4 and −6 mm below 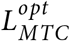) and stimulation durations (35, 65 and 95 ms). CE stimulation was maximal (i.e., *STIM* = 1) during the indicated stimulation duration and fully off elsewhere (see Figure 3C). The combination of four different MTC lengths and three different stimulation durations resulted in twelve different isometric experiments.

#### 2.2.2 *In situ* data

We also used data from an *in situ* experiment on isolated rat m. gastrocnemius medialis, conducted on three male Wistar rats. The experiment included quick-release, step-ramp and isometric experiments, which were similar to those described in Section 2.2.1. For each rat, we performed between 11 and 13 quick-release experiments, 10 and 12 step-ramp experiments and 12 isometric experiments. Below, we provide a brief description of the experimental procedure, which is fully detailed in (Reuvers et al., 2025).

In the experiment, approved by the Committee on the Ethics of Animal Experimentation at the Vrije Universiteit (Permit Number: FBW-AVD11200202114471), rats were first anesthetised with urethane. The hindlimb was then shaved, and the overlying skin and m. biceps femoris were removed. The medial and lateral part of m. gastrocnemius were carefully separated from their surrounding tissue and exposed as much as possible. The rats were placed in the experimental setup with the hindlimb, femur and foot fully fixed. The distal end of the calcaneal tendon was attached to a servomotor (Aurora 309C, Aurora Scientific, Aurora, Canada) using Kevlar thread and aligned to ensure that m. gastrocnemius medialis pulled in its natural direction. All nerves not innervating m. gastrocnemius medialis were severed. M. gastrocnemius medialis was stimulated via a cuff-electrode placed on the sciatic nerve, with the proximal nerves crushed to prevent spinal reflexes. This experimental setup allowed for precise control of m. gastrocnemius medialis length changes and stimulation as well as accurate measurement of m. gastrocnemius medialis force.

### 2.3 Parameter value estimation procedure

The general procedure to estimate the parameter values involved minimising the sum of squared differences between the data and the values based on the estimated parameter values. To distinguish between these, we used subscripts: for example, the value of the SEE force before the quick-release of the data is indicated by 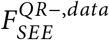 while the value based on the estimated parameter values is indicated by 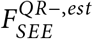.

The data of the quick-release experiments were used to estimate the parameter values of the CE, PEE and SEE force-length relationships. The data of the step-ramp experiments were used to estimate the parameter values of the CE force-velocity relationship. The data of the isometric experiments were used to estimate the parameter values of the excitation dynamics. Below we provide an overview on the methods used to estimate the parameter values. We also offer an open-source toolbox that automates the estimation of contraction and excitation dynamics parameter values.

#### 2.3.1 CE, SEE and PEE force-length parameter value estimation

As explained in the introduction, different combinations of contraction dynamics parameters can yield almost identical mechanical behaviour under isometric conditions (see also Figure 1). For this reason, it is important to estimate SEE stiffness first, in order to discriminate between SEE stiffness on the one hand, and 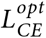 and 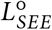 on the other. In our approach, the parameter that scales SEE stiffness (*k*_*SEE*_) is therefore estimated first. This is followed by the estimation of the PEE parameters, as *k*_*SEE*_ is also required for their estimation. Finally, the parameters 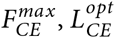, and 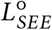 are estimated.

##### Estimation of SEE stiffness

The first step was to estimate SEE stiffness. The model SEE force depends on the parameter values of *k*_*SEE*_ and 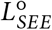, and obviously on SEE length. The problem, however, is that SEE length is unknown in experiments. This makes it challenging to estimate the parameter values concerning the SEE force-length relationship. Fortunately, quick-release experiments provide a way out. During quick-release experiments with a servomotor, the motor quickly shortens MTC length such that there is a rapid decline in SEE force (the ‘quick-release’). Due to the shortness of this timeframe, CE shortening is minimal such that *almost all* MTC shortening is taken up by SEE. Often, in experimental studies, it is assumed that *all* MTC shortening can be attributed to SEE shortening. In reality, this assumption leads to an overestimation of SEE shortening, as CE also shortens during this short timeframe. We introduced a method to correct for this overestimation of SEE shortening (see Section 2.3.4) and estimated the parameter values considering both methods: without and with correcting for CE shortening during the quick-release.

Obtaining the SEE length change due to the quick-release 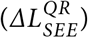 is a crucial step. This is because SEE length after the quick-release 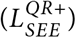 can then be expressed as SEE length before the quick-release 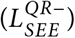 plus the change in SEE length due to the quick-release: 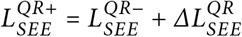. This allowed us to rewrite Equation 2 to estimate SEE force immediately before 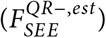 and after 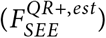 the quick-release. This yielded:

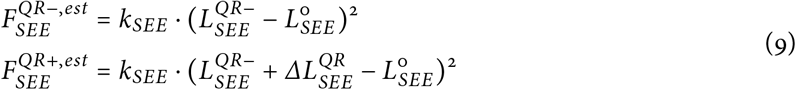

Now, there are two unknown parameters (i.e., *k*_*SEE*_ and 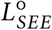), while also SEE length before the quick-release is also unknown for each quick-release experiment. Consequently, there are more unknowns than equations. To address this issue, we replaced 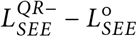 with a temporary parameter *c*_*SEE*_. This yielded:

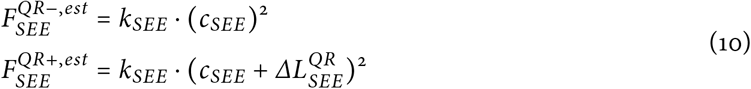

As *k*_*SEE*_ scales SEE stiffness, its value is constant across all quick-release experiments. In contrast, *c*_*SEE*_ depends on the parameter 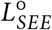 and SEE length before the quick-release, which differs among each quick-release experiment. Consequently, *c*_*SEE*_ should be determined for each quick-release experiment individually. Visually, *c*_*SEE*_ determines the shift along the x-axis on the SEE force-length relationship, while *k*_*SEE*_ scales SEE stiffness and consequently SEE force (see Figure 4A). The value of *k*_*SEE*_ (and *c*_*SEE*_ for each quick-release experiment) were found by minimising the sum of squared differences between the data and estimated SEE force before 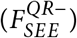 and after 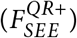 the quick-release.

**Figure 4:**
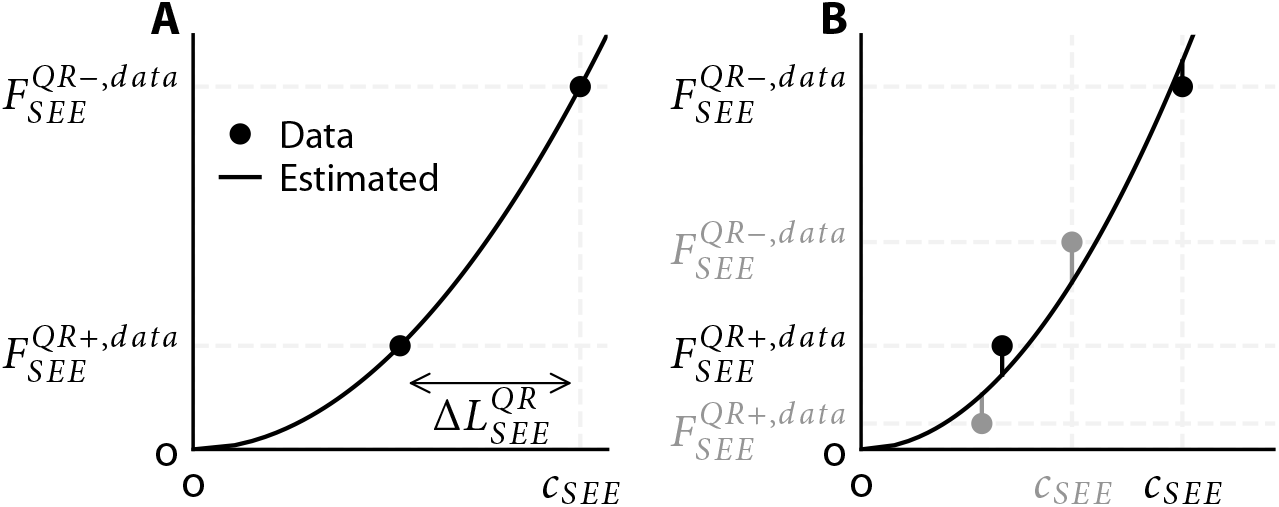
Graphical illustration of SEE stiffness parameter estimation. For each quick-release experiment, SEE force before 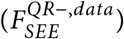 and after 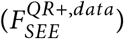 the quick-release was obtained from the data, as well as the corresponding decrease in SEE length 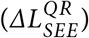. Since SEE length prior to the quick-release is typically unknown, a temporary parameter (*c*_*SEE*_) was introduced for each experiment to represent the difference between SEE length before the quick-release and SEE slack length. A) A single quick-release experiment yields two unknowns: *c*_*SEE*_ and a parameter that scales SEE stiffness (*k*_*SEE*_). B) Running multiple quick-release experiments yields n+1 unknowns: n values of *c*_*SEE*_ (one for each experiment) and the parameter scaling SEE stiffness. Here, two quick-release experiments are illustrated (one indicated with black dots, the other with grey dots), while the estimated SEE force-length relationship is depicted with the black line.

##### Estimation of PEE parameter values

The second step was to estimate PEE stiffness. The model PEE force depends on the parameter values of *k*_*PEE*_ and 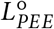, and obviously on PEE length. In experiments, SEE force is measured, while PEE force is required to estimate PEE stiffness. As such, experimental data should be used in which CE force is negligible such that PEE force is approximately equal to SEE force. At the beginning of the quick-release experiments, the active state is so low that CE force is negligible. We selected an interval of 10 ms in which the SEE force was minimal (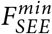, orange dot in Figure 3A).

In experiments, PEE length is unknown. This problem can be addressed as follows. First, PEE length equals MTC length minus SEE length: *L*_*PEE*_ = *L*_*MTC*_ − *L*_*SEE*_ (Equation 1). Second, SEE length is the sum of SEE slack length and SEE elongation 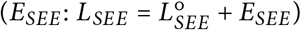. This allowed us to rewrite Equation 2 to estimate PEE force 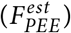, yielding:

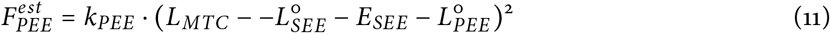

Now, there are two unknown parameter values (i.e., *k*_*PEE*_ and 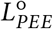), while also SEE elongation is unknown for every quick-release experiment. Consequently, there are more unknowns than equations. To address this, we replaced 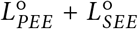 with a temporary parameter *c*_*PEE*_. Subsequently, SEE elongation was computed as 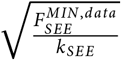, given the estimated SEE stiffness scaling factor in the previous step. This yielded:

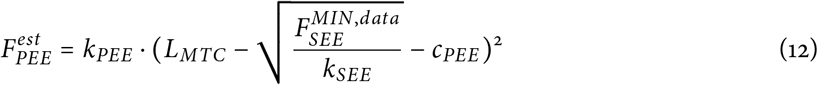

As *k*_*PEE*_ only scales PEE stiffness, its value is constant across all quick-release experiments. Similarly, *c*_*PEE*_ is the sum of 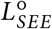 and 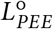 and should therefore be constant for each quick-release experiment. The values of *k*_*PEE*_ and *c*_*PEE*_ were then computed by minimising the sum of squared differences between 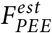 and 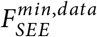.

##### Estimation of 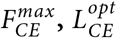 and 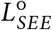

The third step was to estimate 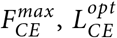 and 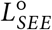. Before the quick-release, MTC is isometrically delivering force. We used the SEE force at this instant (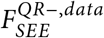, black dot in Figure 3A) and the corresponding MTC length to obtain the MTC force-length relationship from the data. We then estimated maximal isometric CE force 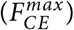, CE optimum length 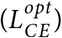 and SEE slack length 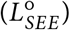 by minimising the sum of squared differences between the data and estimated MTC force-length relationship. Lastly, 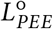 was computed by subtracting 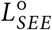 from *c*_*PEE*_.

#### 2.3.2 CE force-velocity parameter value estimation

The values of the CE force–velocity relationship parameters *a* and *b* were estimated from data obtained during the plateau phase of SEE force in step-ramp experiments. This approach was chosen because SEE length is constant when SEE force is constant. Consequently, CE velocity equals MTC velocity under these conditions. Following this argument, we first identified a 10 ms interval in which SEE force changed the least. Second, we derived CE length and CE force as functions of time using Equation 1 and Equation 2. Third, we calculated the CE velocity as the time-derivative of CE length. Finally, we averaged CE force and CE velocity over the 10 ms interval. This procedure yielded the data of the CE force-velocity relationship.

Now, the model CE force-velocity relationship can be fit to that of the data to obtain value of *a* and *b*. However, employing a method that simply minimises the sum of ordinary least squares differences would not be appropriate because in experiments there is uncertainty in both measured CE force and CE velocity. Instead, we used a total least squares method that minimised the distance between the modelled CE force 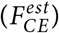 and CE velocity 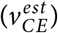 (now called: model values) and those of the data (see also Figure 5). To do this, we had to find the nearest model values to the CE force of the data 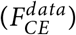 and CE velocity of the data 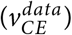 (now called: datapoints). We used the following observations: 1) the model value has to satisfy Equation 4 and 2) the derivative of 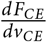 of the model value should be perpendicular to the line from the datapoint to the model value (see Figure 5). Hence, the model value could be found by solving Equation 4 and Equation 13:

**Figure 5:**
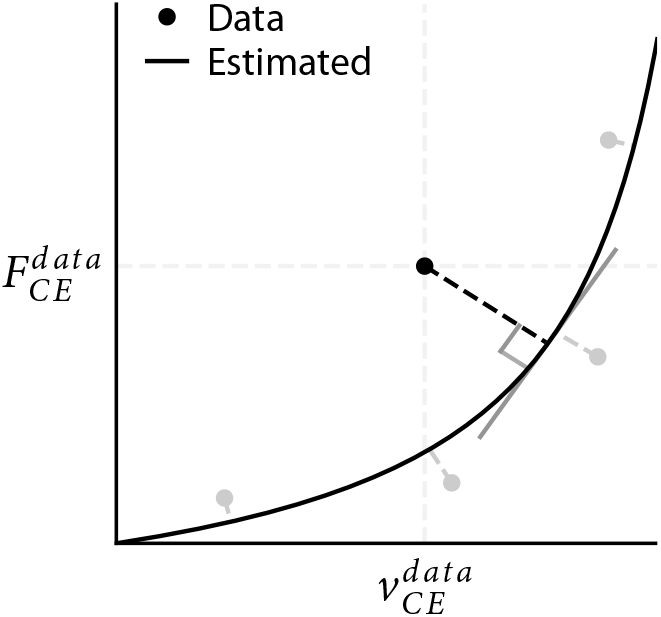
Graphical illustration of the CE force-velocity parameter estimation using total least squares. For each step-ramp experiment, CE force and CE velocity was obtained during the interval in which SEE force changed the least. The nearest point on the CE force-velocity relationship was identified based on two criteria: it should satisfy Equation 4, and the line from the datapoint to the CE force-velocity relationship should be perpendicular the CE force-velocity relationship. This was done for all step-ramp experiments (other datapoints are depicted in grey), while the estimated CE force-velocity relationshp is depicted with the black line.

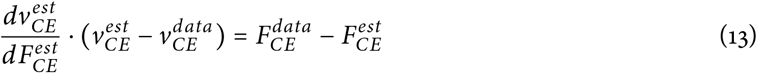

The following cost-function was then minimised to find the parameter values of *a* and *b*:

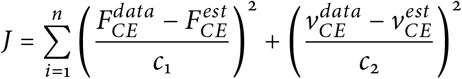

*c*_1_ and *c*_2_ denote scaling factors such that both terms of the cost-function are more or less equally weighted, which was done by setting *c*_1_ equal to the maximal range in CE force of the data and setting *c*_2_ equal to the maximal range of CE velocity of the data.

#### 2.3.3 Excitation dynamics parameter value estimation

We estimated the parameters values of *τ*_*act*_ and *τ*_*deact*_ based on isometric experiments. The choice to estimate only these parameters of the excitation dynamics was made because it is generally challenging to discriminate between parameter values of both the calcium dynamics (Equation 7) and the *q* − [*Ca*^2+^] relation (Equation 8) based on mechanical measurements outlined above. The parameter values of the *q* − [*Ca*^2+^] relation were set to those used in simulating the isometric experiments. Consequently, these parameters matched their actual values for the simulated data, but obviously were suboptimal for the *in situ* data. To estimate the parameter values of *τ*_*act*_ and *τ*_*deact*_, we minimised the sum of squared differences between the SEE force of the data and the estimated SEE force based on the parameter values. For this, an interval of the data was used starting from maximal CE stimulation to 0.1 s after CE stimulation ceased off.

#### 2.3.4 Parameter estimation: traditional method & improved method

To estimate the SEE stiffness it is generally assumed that the quick-release is so fast that all MTC shortening is taken up by SEE. As A.V. Hill already acknowledged in (1950), this is not the case in reality as CE also shortens. Although this CE shortening might be small, SEE is in general stiff and therefore small errors in SEE length may result in large errors in predicted SEE forces and thus in estimated SEE stiffness. We used two different methods to estimate the parameter values: 1) the traditional method and the improved method. In the traditional method, we followed the procedure outlined in section Section 2.3.1 and Section 2.3.2. In the improved method, we also followed the procedure outlined in Section 2.3.1 and Section 2.3.2 after which we incorporated an additional procedure to correct for CE shortening due to the quick-release. This additional procedure was as follows: 1) We calculated CE length just before the (step) change in MTC length occurred using Equation 1, 2, 3 and 8, assuming that CE velocity was 0 and that *γ* equalled 1. 2) We performed a (short) simulation from the time just before the (step) change in MTC length to just after the (step) change in MTC length. 3) We computed the change in CE length over this time interval as the average CE length slightly before and slightly after the (step) change in MTC length. 4) We subtracted the change in CE length from the change in MTC length to obtain the change in SEE length. In addition to correcting for CE length change during the quick-release, we incorporated a step to obtain better estimates of the actual PEE force at the time instance of minimum SEE force (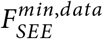; see Figure 3). We computed CE force at this time instance using Equation 3 and 8, assuming that *γ* equalled its minimum value. This CE force was then subtracted from 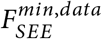.

The improved method leads to a different (i.e., higher) SEE stiffness, which affect the estimation of the parameter values of the CE, PEE and SEE force-length relationships (see Section 2.3.1). These estimated parameter values were then used to again estimate the parameter values of the CE force-velocity relationship (see section Section 2.3.2). As the parameter values of the CE force-velocity relationship slightly changed, the estimate of CE length change due to the quick-release also changed and therefore estimated SEE stiffness changed. We found that this process always converged to a stable solution and therefore we used an iterative process until the change in all parameter values was less than 0.1% (see Figure S2).

### 2.4 Sensitivity analysis

#### 2.4.1 Monte Carlo simulations

In experiments on isolated MTC, the MTC force-length relationship can shift due to irreversible damage of SEE (Aubert et al., 1951). We observed this phenomenon in an experiment involving isolated rat m. gastrocnemius medialis, where the MTC force-length relationship shifted by about 1 mm 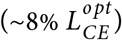 over a period of approximately 8 hours (Reuvers et al., unpublished observation). To assess the influence of these shifts on the parameter value estimation, we performed Monte Carlo simulations. We assumed that the observed shifts of the MTC force-length relationship were caused solely by a decrease in SEE stiffness. Accordingly, we decreased SEE stiffness to cause random shifts of the MTC force-length relationship between 0 and 1 mm. This way, we simulated data as if they were collected at random intervals throughout an 8-hour period. Importantly, changes in SEE stiffness affect the step size and constant MTC velocity ramp required for a more or less a constant SEE force at the beginning of the ramp in step-ramp experiments. Therefore, we adjusted the step size and constant MTC velocity ramp for each simulated step-ramp trial individually. Parameter values of each MTC were then estimated using the ‘perturbed’ data using the improved method (see Section 2.3.4). This process was repeated 50 times for each MTC, allowing us to evaluate both the mean and the spread of the estimated parameter values.

#### 2.4.2 Interdependency of parameter values

The parameter estimation procedure detailed above follows a fixed order in which certain parameters are estimated before others. For instance, *k*_*SEE*_ is estimated first and therefore affects the estimation of 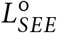 and 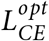. This sequence creates an interdependency among the parameter values, which has been identified as a potential cause for substantial variation in parameter values among individuals, even within the same muscle type (Lemaire et al., 2016). To investigate this interdependency, we systematically changed parameter values of our model. Specifically, we increased and decreased each parameter value by 5% relative to the values obtained with the improved method. After this, we re-estimated all other parameter values. This approach allowed us to assess the influence of changes in one parameter on the estimation of others.

## 3 Results

### 3.1 Evaluating parameter value estimation accuracy - simulated data

#### 3.1.1 Traditional method versus Improved method

The traditional method did not correct for CE shortening during quick-release experiments. Consequently, SEE shortening due to the quick-release was overestimated by 24% on average. This overestimation of SEE shortening caused a decrease in *k*_*SEE*_ (and thus SEE stiffness) of 34% on average. The estimation of the activation time constant was most affected by the underestimated SEE stiffness, resulting in an underestimation of 19% on average. All other estimated parameter values were within 9% of their actual values (Table 1). These findings show the influence of CE shortening — even within a very brief 10 ms interval — on the parameter value estimation of both the contraction dynamics and excitation dynamics.

**Table 1:** Percentage differences between estimated and actual MTC parameter values.

|  | *Traditional method* |  |  | *Improved method* |  |  | *Monte Carlo* |  |  |
| --- | --- | --- | --- | --- | --- | --- | --- | --- | --- |
|  | GM1 | GM2 | GM3 | GM1 | GM2 | GM3 | GM1 | GM2 | GM3 |
| $a$ | 1 | 4 | 2 | 1 | 3 | 1 | $1 \pm 2$ | $6 \pm 2$ | $2 \pm 3$ |
| $b$ | -0 | 2 | -1 | 1 | 2 | 1 | $1 \pm 2$ | $5 \pm 2$ | $2 \pm 3$ |
| $F_{CE}^{max}$ | 0 | -1 | 0 | -0 | -1 | -0 | $-0 \pm 1$ | $-2 \pm 1$ | $-1 \pm 1$ |
| $k_{PEE}$ | 3 | -9 | 9 | -0 | 0 | -0 | $3 \pm 17$ | $3 \pm 15$ | $7 \pm 31$ |
| $k_{SEE}$ | -39 | -27 | -37 | 0 | -1 | 0 | $-37 \pm 6$ | $-36 \pm 6$ | $-35 \pm 6$ |
| $L_{CE}^{opt}$ | -1 | -2 | -3 | -0 | 0 | -0 | $0 \pm 3$ | $1 \pm 3$ | $0 \pm 3$ |
| $L_{PEE}^o$ | 0 | -1 | -0 | -0 | 0 | -0 | $0 \pm 3$ | $1 \pm 2$ | $0 \pm 2$ |
| $L_{SEE}^o$ | -1 | -0 | -0 | 0 | -0 | 0 | $-0 \pm 1$ | $-0 \pm 1$ | $-0 \pm 1$ |
| $\tau_{act}$ | -21 | -16 | -21 | -0 | -1 | -0 | $1 \pm 3$ | $1 \pm 4$ | $1 \pm 4$ |
| $\tau_{deact}$ | -5 | -4 | -5 | 0 | 0 | 0 | $0 \pm 1$ | $0 \pm 1$ | $0 \pm 1$ |

**Table 2:** Interdependency of the estimated MTC parameter values. Each entry shows the percentage change in the row parameter resulting from a 5% change in the column parameter. All values are expressed as percentage changes.

| | $a$ | $b$ | $F_{CE}^{max}$ | $k_{SEE}$ | $L_{CE}^{opt}$ | $L_{SEE}^o$ |
| --- | --- | --- | --- | --- | --- | --- |
| $a$ | - | $10.2 \pm 1.1$ | $-14.1 \pm 1.7$ | $-0.1 \pm 0.0$ | $3.4 \pm 0.4$ | $-10.3 \pm 1.5$ |
| $b$ | $2.3 \pm 0.2$ | - | $-14.2 \pm 2.0$ | $0.1 \pm 0.0$ | $3.8 \pm 0.5$ | $-10.7 \pm 1.3$ |
| $F_{CE}^{max}$ | $-0.0 \pm 0.0$ | $-0.0 \pm 0.0$ | - | $-0.1 \pm 0.0$ | $-1.2 \pm 0.2$ | $3.6 \pm 0.8$ |
| $k_{PEE}$ | $-0.2 \pm 0.1$ | $-0.4 \pm 0.1$ | $2.2 \pm 0.5$ | $-1.3 \pm 0.4$ | $-0.5 \pm 0.1$ | $1.4 \pm 0.3$ |
| $k_{SEE}$ | $0.6 \pm 0.2$ | $1.5 \pm 0.3$ | $-6.5 \pm 2.1$ | - | $1.6 \pm 0.4$ | $-4.2 \pm 1.1$ |
| $L_{CE}^{opt}$ | $0.0 \pm 0.0$ | $0.0 \pm 0.0$ | $-5.3 \pm 0.6$ | $0.2 \pm 0.1$ | - | $-14.0 \pm 1.3$ |
| $L_{PEE}^o$ | $0.0 \pm 0.0$ | $0.0 \pm 0.0$ | $0.0 \pm 0.0$ | $0.0 \pm 0.0$ | $0.0 \pm 0.0$ | $-10.5 \pm 0.7$ |
| $L_{SEE}^o$ | $0.0 \pm 0.0$ | $0.0 \pm 0.0$ | $1.8 \pm 0.3$ | $0.1 \pm 0.0$ | $-1.7 \pm 0.1$ | - |
| $\tau_{act}$ | $0.6 \pm 0.2$ | $1.6 \pm 0.2$ | $-3.3 \pm 1.1$ | $2.3 \pm 0.2$ | $1.1 \pm 0.6$ | $-3.5 \pm 2.6$ |
| $\tau_{deact}$ | $0.2 \pm 0.0$ | $0.5 \pm 0.0$ | $-2.9 \pm 0.2$ | $0.6 \pm 0.0$ | $1.2 \pm 0.0$ | $-2.8 \pm 0.3$ |

In contrast, the improved method corrected for CE shortening during the quick-release experiments. The correction substantially reduced the overestimation of SEE shortening, yielding an accurate estimate of *k*_*SEE*_. As a result, all estimated parameter values deviated by no more than 3% from their actual values (Table 1). These results demonstrate that accounting for CE shortening - even over a brief 10 ms interval - substantially improves the accuracy of the parameter value estimation.

For interested readers, we discuss below the specific factors contributing to the differences between the actual contraction and excitation dynamics parameter values and those estimated based on the traditional method.

#### 3.1.2 Understanding errors in parameter value estimation

##### Estimation of SEE stiffness

The SEE stiffness parameter (*k*_*SEE*_) was underestimated by 39%, 27% and 37% for GM1, GM2 and GM3, respectively (see Table 1). This overestimation directly resulted from the overestimation of SEE shortening due to the quick-release, which was overestimated by 28±2%, 17±2% and 26±2% for GM1, GM2 and GM3, respectively, averaged across all quick-release experiments. In absolute terms, CE shortened only 43±3, 29±2 and 41±2 µm over the 11 ms interval between the time points at which SEE force and SEE length were sampled (i.e., immediately before and after the quick-release). The amount of CE shortening was smallest in GM2 because GM2 was a slower muscle than GM1 and GM3. Consequently, the overestimation of SEE shortening was also smallest in GM2, which in turn led to the smallest underestimation of *k*_*SEE*_. All in all, this shows that even very small amounts of CE shortening has substantial influence on the estimation of SEE stiffness.

Since SEE was very stiff, the overestimation in SEE elongation at maximal isometric CE force was only 0.50, 0.32 and 0.49 mm for GM1, GM2 and GM3, respectively. This is an important finding because the SEE stiffness is the first parameter in the estimation process and therefore affects all subsequent estimated parameter values. Therefore, since the overestimation of SEE elongation (in mm) at maximal isometric CE force was only small, the impact on the estimation of the other contraction dynamics parameter values was also small.

Lastly, it should be noted that the correlation coefficient between the data SEE force-length relationship and the one based from the estimated parameters is not an adequate measure of how accurately SEE stiffness is captured. Obviously, a correlation coefficient is not sensitive to the overestimation of SEE shortening due to the quick-release. Consequently, high correlation coefficients (*R*^2^ > 0.99) can still be observed even when SEE stiffness is substantially underestimated. To further investigate this issue, we simulated the quick-release experiments after obtaining all contraction and excitation dynamics parameter values. The simulations clearly showed a slower rise in SEE force after the quick-release in comparison with the experimental data (Figure 6A), indicating that SEE stiffness was underestimated. These results suggest that the rise in SEE force is a better measure of SEE stiffness estimation accuracy.

**Figure 6:**
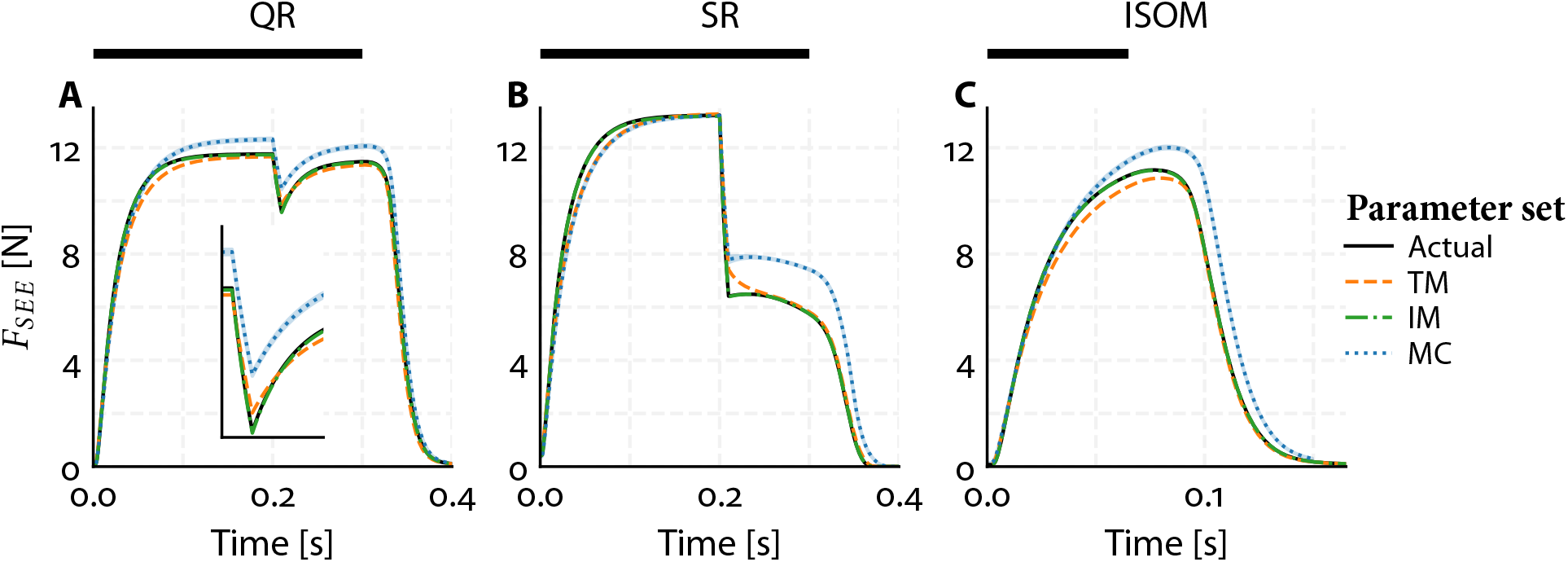
Representative example of SEE force over time during a quick-release (A), step-ramp (B) and isometric experiment (C). The inset in (A) depicts the SEE force over time around the quick-release. The SEE force over time is depicted for four sets of parameter values: 1) the actual values (i.e., literature-obtained; black solid line), those obtained with the traditional method (TM; orange dashed line), those obtained with the improved method (IM; green dashed-dotted line) and those resulting from the Monte Carlo Simulations (MC; blue dotted line, with shaded 95% confidence interval).

**Figure 7:**
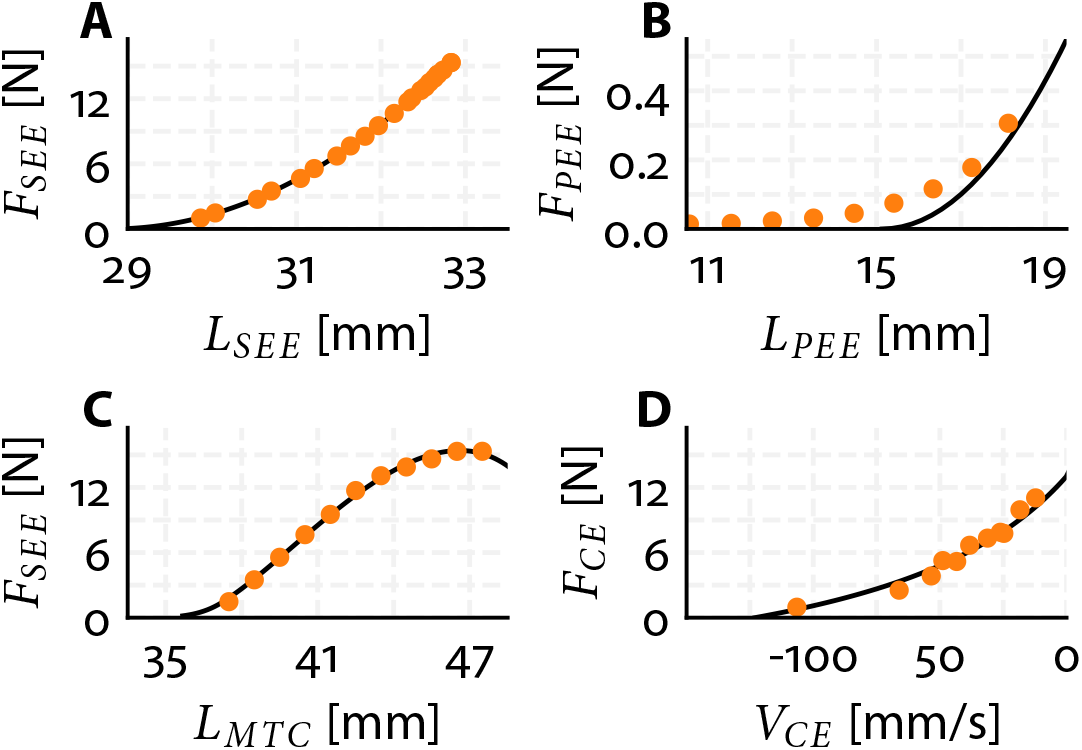
Representative example of experimental *in situ* data of rat 1 for the SEE force-length relationship (A), PEE force-length relationship (B), MTC force-length relationship (C) and the CE force-velocity relationship (D). The orange dots depicts the experimental *in situ* data and the solid black line depicts the model fit.

##### Estimation of PEE parameter values

The PEE stiffness parameter (*k*_*PEE*_) was 2.8% higher, 8.6% lower and 8.7% higher than the actual parameter value for GM1, GM2 and GM3, respectively (see Table 1). PEE slack length 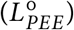 values were 0.29% higher, 1.4% lower and 0.048% lower than the actual values for GM1, GM2 and GM3, respectively (see Table 1). The available data to estimate 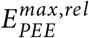 and 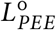 was limited as in most quick-release experiments PEE length was below 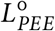. In fact, only 4, 4 and 3 datapoints were used to estimate the PEE force-length relationship for GM1, GM2 and GM3, respectively. Notwithstanding, the estimated PEE parameter values were within 8.7% of their actual value.

##### Estimation of 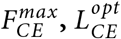 and 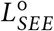

The estimated values of 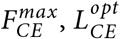, and 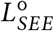 were within 2.6% of their actual values (see Table 1). The deviations were all due to the underestimation of *k*_*SEE*_, which can be explained in detail as follows. First, the width of the model’s MTC force-length relationship is mainly determined by *k*_*SEE*_ and 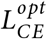. The lower *k*_*SEE*_, the wider the MTC force-length relationship; the shorter 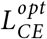, the narrower the relationship. Consequently, underestimating *k*_*SEE*_ caused a underestimation of 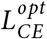. Second, a lower 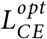 results in a lower 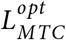. This led to 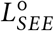 being overestimated such that 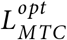 of the data and model were aligned. Third, a lower *k*_*SEE*_, lower 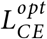 and higher 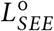 results in a narrower MTC force-length relationship (see Figure 1A). This resulted in an increase 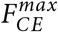 in order to minimise the root mean squared distance between estimated MTC force-length relationship and that of the data. Therefore, the underestimation of SEE stiffness resulted in a small overestimation of 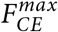, and 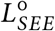, and a small underestimation of 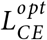.

##### Estimation of CE force-velocity parameter values

The estimated value of *a* was overestimated by up to 3.8%, while *b* was within 1.6% of its actual value (see Table 1). The slight overestimation of *a* resulted from the estimation of CE force and CE velocity over a 10 ms interval in which the SEE force changed the least (see Figure 3B). At high shortening velocities, CE length decreased so rapidly that — even over a short interval of 10 ms — SEE force changed. Consequently, assuming that CE velocity equalled MTC velocity led to an overestimation of CE velocity. These findings show that it is crucial in experiments to find a combination of step sizes and constant MTC velocity ramps resulting in a nearly constant SEE force. While this is experimentally challenging, achieving this would allow for accurate estimation of the CE force-velocity relationship.

##### Estimation of excitation dynamics parameter values

*τ*_*act*_ and *τ*_*deact*_ were underestimated across all muscles. For GM1, GM2, and GM3, *τ*_*act*_ was underestimated by 21%, 16% and 21%, respectively, while *τ*_*deact*_ was underestimated by 4.9%, 3.6% and 5.1%, respectively (see Table 1). This underestimation was primarily due to the underestimation of SEE stiffness. In short, the underestimation of SEE stiffness caused an overestimation of SEE lengthening velocity, which in turn lead to an overestimation of CE shortening velocity. This reduced CE force via the CE force-velocity relationship and therefore a slower increase in SEE force. Consequently, *τ*_*act*_ was overestimated, leading to faster increase in SEE force. The underestimation of *τ*_*act*_ also resulted in an overestimation of the relative [*Ca*^2+^] at the start of the deactivation, causing a decrease in *τ*_*deact*_ (leading to a faster decrease in SEE force). This finding highlights the influence of the contraction dynamics on the estimation of the excitation dynamics parameter values.

To illustrate the influence of the contraction dynamics on the estimation of the excitation dynamics parameter values, we also fitted *τ*_*act*_ and *τ*_*deact*_ to the data of the quick-release and step-ramp experiments. Using the quick-release data, *τ*_*act*_ was underestimated by 38% on average and based on step-ramp data *τ*_*act*_ was underestimated by 64% on average. In turn, *τ*_*deact*_ was underestimated by only 1% on average for both data sets. These results show that quick-release and step-ramp experiments are not suitable for accurately estimating *τ*_*act*_ due to substantial CE length changes that occur in these experiments. To minimise the influence of contraction dynamics and improve the estimation of *τ*_*act*_, experiments should be performed in which CE length changes are minimal. Isometric experiments are best suited for this purpose.

### 3.2 Sensitivity analysis

#### 3.2.1 Monte Carlo simulations

Monte Carlo simulations were used to examine how shifts in the MTC force–length relationship caused by a decrease in SEE stiffness (e.g., due to irreversible SEE damage in experiments) affect the accuracy of the estimated contraction and excitation dynamics parameter values. The induced shifts of the MTC force-length relationship were between 0–1 mm, and therefore it was no surprise that SEE stiffness decreased by 36% on average — which corresponds to a 0.5 mm shift.

All other estimated contraction dynamics parameter values were within 6.3% on average of their actual value (Table 1). The variance in the estimated contraction dynamics parameter values was below 31% for all parameters except for the PEE stiffness scaling parameter. The PEE stiffness scaling parameter showed substantially higher variance (with a standard deviation up to 31%), because it was based on four or fewer quick-release trials, making it more sensitive to perturbations in the data. Regarding the excitation dynamics, the influence on the average time constants was minimal (within 0.71%), but affected the variance in the estimated values of the activation time constant (with a standard deviation of 4% on average). These findings indicate that an average decrease in SEE stiffness of 36% has a much smaller effect on the estimated parameter values of the contraction and excitation dynamics, even at the level of an individual muscle.

#### 3.2.2 Interdependency of parameter values

We investigated the interdependency of the estimated parameter values by adjusting each parameter value by 5% and re-estimating all other parameter values. 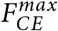 and 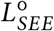 were the parameters that had most effect on the estimates of the others.

First, 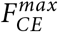 affected the estimation of parameter *a* and *b* of the CE force-velocity relationship. Under-estimating 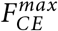 leads to an underestimation of CE force at 0 velocity. Due to the formulation of the CE force-velocity relationship, the curve, by definition, crosses the point at *v*_*CE*_ *=* 0 and 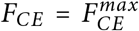. Consequently, the best fit to the data with an underestimated 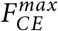 is a much flatter CE force-velocity relationship, which is realised by an increase in *a* and a decrease in *b*. This flatter CE force-velocity relationship affects the estimation of all other contraction dynamics parameter values because the CE force-velocity relationship is used to estimate the CE shortening due to the quick-release (see Section 2.3.4). As a result, *k*_*SEE*_ and 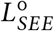 decreased due to underestimating 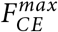, whereas 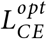 increased. In summary, 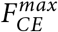 substantially influenced the estimation of all contraction dynamics parameter values mainly by its effect on the CE force-velocity relationship.

Second, 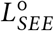 affected the estimation of 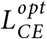 and 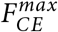. As explained earlier, when 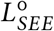 is overestimated, 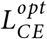 is underestimated, such that 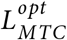 remains more or less unaffected. Underestimating 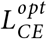 causes a narrower MTC force-length relationship, leading to an overestimation of 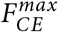 to preserve a good fit between the data and the model. As previously discussed, overestimating 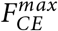, in turn, resulted in an underestimation of parameters *a* and *b*. Thus, 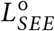 substantially influenced the estimation of all contraction dynamics parameter values mainly by its effect on the MTC force-length relationship.

Taken together, the interdependence of 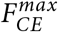 and 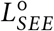 with other parameter values underscores the need for precise estimation of these key parameters. In this regard, it is reassuring that these two parameters were found to be robust for perturbations in the experimental data.

### 3.3 Evaluating model predictions - *in situ* data

We estimated contraction and excitation dynamics parameter values from *in situ* data of rat m. gastrocnemius medialis (Table S2). The improved method yielded a 67% higher SEE stiffness compared to the traditional method. As explained earlier, higher SEE stiffness also affects the estimation of all other parameter values. The most noticeable changes were longer time constants for both the activation (*τ*_*act*_) and deactivation (*τ*_*deact*_) dynamics, with increases of 61% and 16% on average, respectively. Another noticeable change was in CE optimum length, which was about 11% higher using the improved method compared to the traditional method.

To assess model predictions, we re-ran quick-release, step-ramp and isometric experiments, as well as stretch-shortening cycles with experimentally measured MTC length and CE stimulation over time as inputs to the model. We compared the model predictions using parameter sets obtained from both methods. For all experiments and rats, the average difference between experimental data and model predictions was 30% smaller on average using parameters from the improved method (Figure 8; Table S3). Overall, the improved method substantially enhanced the predictions derived by the Hill-type MTC model compared to the traditional method.

**Figure 8:**
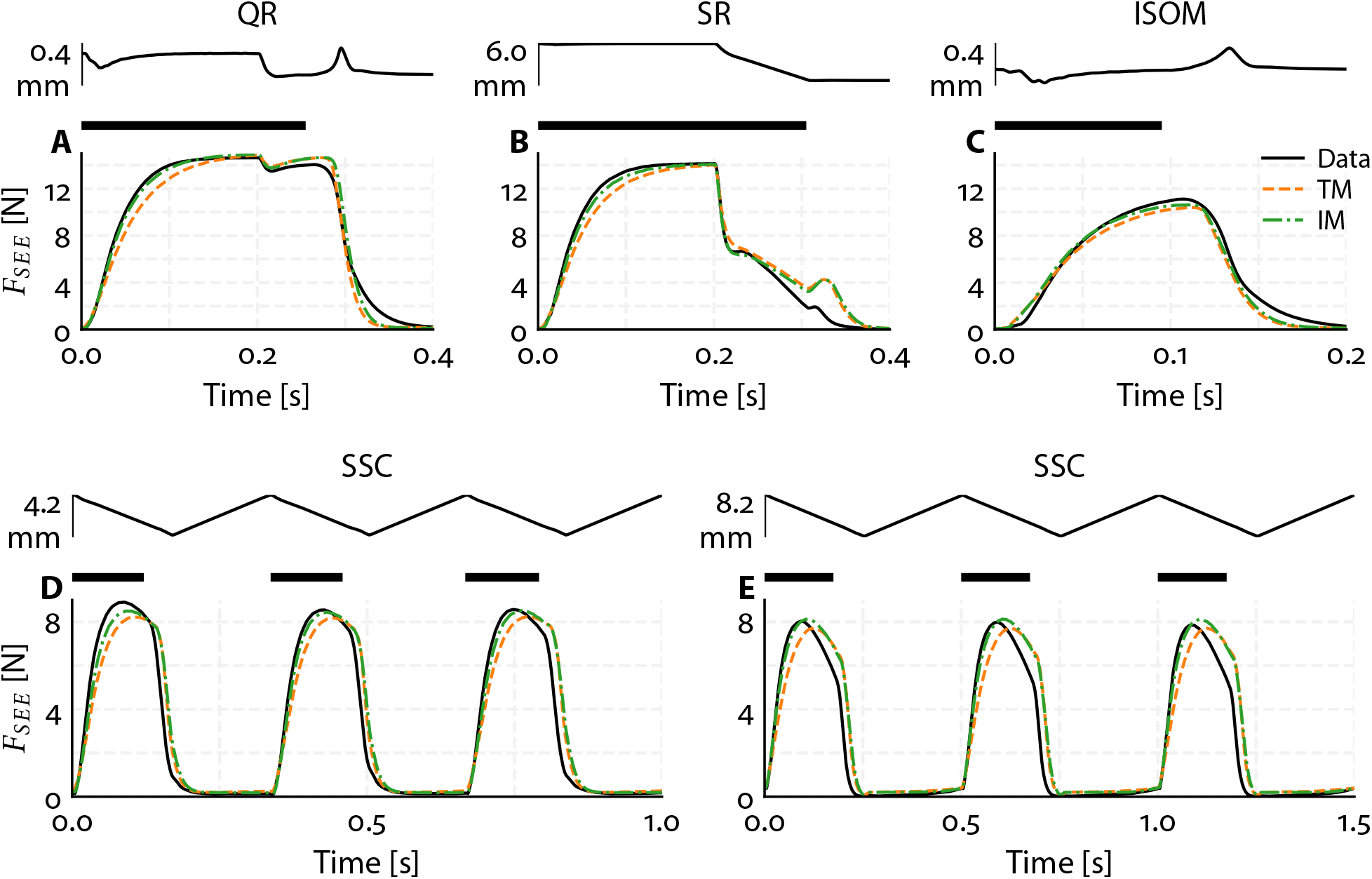
Representative example of experimental *in situ* data and simulation results of rat 1 for SEE force over time during a quick-release experiment (A), step-ramp experiment (B), isometric experiment (C) and two stretch-shortening cycles (D & E). The simulation results were obtained by re-simulating the experimental protocol with the MTC length and CE stimulation from the experimental data as input and by using either the parameter set obtained with the traditional method (TM; orange dashed line) or the improved method (IM; green dash-dotted line). For each panel, the top plot represents MTC length over time, with the bar indicating its range in mm. CE stimulation is maximal during the periods indicated by the black bars and is ‘off’ elsewhere.

## 4 Discussion

The aim of this study was to evaluate the accuracy of estimating contraction and excitation dynamics parameter values on the basis of data collected in commonly used experimental protocols. In real experiments, the actual parameter values are unknown, making it impossible to directly assess estimation accuracy. In this study, we took a different approach: using a Hill-type MTC model with parameter values obtained from literature we generated synthetic ‘data’ by simulating quick-release, step-ramp, and isometric experiments. We then estimated all important contraction and excitation dynamics parameter values by using the synthetic data. Since the actual parameter values were known in this case, we could assess how accurately they were retrieved. Two different estimation methods were compared. The first was the traditional method, commonly used in the literature, which does not account for muscle fibre shortening during quick-release experiments. The second was an improved method that includes a correction for muscle fibre shortening during the quick release. Both methods developed in this study—designed to estimate contraction and excitation dynamics parameters from quick-release, step-ramp, and isometric experiments using servomotors—are made available as an open-source toolbox. In the remainder of this paper, we will discuss 1) the difference between the two methods; 2) the robustness of the improved method and 3) the implications for muscle modelling.

In quick-release experiments using servomotors, muscle fibres shorten, even within the short duration of the release. Obviously, the extent of muscle fibre shortening depends on the duration of the quick-release. Consequently, the longer the quick-release, the more important it becomes to account for muscle fibre shortening when estimating SEE stiffness (see Figure S3). For a typical quick-release duration of 10 ms, we found that muscle fibre shortening was nearly 25% of MTC shortening (and thus 33% of SEE shortening), resulting in an underestimation of SEE stiffness by approximately 35%. This finding is important, especially in studies aiming to estimate SEE energy storage from SEE force or length, since energy storage estimates scale linearly with SEE stiffness. Although SEE was substantially underestimated with the traditional method, this had minimal effect on the estimated parameter values of the PEE and CE force-length relationships and the CE force-velocity relationship, but impacted the estimated excitation dynamics parameter values. The improved method, in turn, yielded SEE stiffness values close to the actual ones and substantially enhanced the estimates of the other parameter values.

Our sensitivity analysis revealed substantial interdependencies between certain parameters. Particularly, we observed high sensitivity of estimated parameter values to variations in the maximal isometric CE force 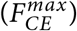 and the SEE slack length 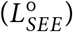 parameter values. Despite this sensitivity, it is important to note that these parameters were accurately estimated even from perturbed experimental data. This was true in general: contraction and excitation dynamics parameter values were quite robust against shifts in the MTC force-length relationship of the simulated data according to the Monte Carlo simulations. These findings indicate the robustness of the improved method.

We applied both the traditional and the improved method to *in situ* data from quick-release, step-ramp, and isometric experiments on rat m. gastrocnemius medialis. The improved method yielded substantially higher estimates of SEE stiffness compared to the traditional method (67% on average). Using parameter sets from both methods, we simulated the quick-release, step-ramp, and isometric experiments, as well as independent stretch-shortening cycles, all with experimentally obtained MTC length and CE stimulation over time as input to the model. With parameter estimates obtained with the traditional method, the root mean squared difference between predicted and experimentally measured force was about 5.6% of maximal CE force, averaged across all rats and experiments; with parameter estimates obtained with the improved method this was only 3.9%. The small difference between predicted and experimentally observed force over time with parameters from the improved method suggests that muscle force can be accurately predicted with a Hill-type MTC model across a wide variety of contractions. This is a remarkable finding for two reasons. First, the Hill-type MTC model is obviously a simplification of real muscle, abstracting the muscle belly, aponeurosis, and tendon into the CE, SEE, and PEE components, and thus does not account for certain complexities (e.g., non-constant pennation angle, muscle inhomogeneities, etc.). Second, the Hill-type MTC model used in this study consists of 26 parameters, of which only 10 values were estimated. This means that all other parameters were left suboptimal. Despite these simplifications, the model was able to accurately predict muscle behaviour, demonstrating that a relatively simple model with a limited number of estimated parameter values can still capture important aspects of muscle dynamics.

In conclusion, our results demonstrate that the improved parameter estimation method provides accurate and robust estimates of contraction and excitation dynamics properties, offering a reliable approach for muscle property estimation in both biomechanical and muscle physiology research. The approach that we designed in this study is made available as an open-source toolbox (https://github.com/edwinreuvers/mp-estimator).

## Acknowledgments

The authors thank Maarten F. Bobbert for helpful comments on a draft of the paper. The authors also thank Koen K. Lemaire for insightful discussions.

## Data and resource availability

All data, code, and materials used in this study are openly available:

- **GitHub repository:** All raw data, processed data, and analysis code are hosted on GitHub at https://github.com/edwinreuvers/acc-mp-estimation.
- **Reproducible analysis website:** Full analysis pipeline — including data, analysis code, and figure/table generation — is available at https://edwinreuvers.github.io/publications/acc-mp-estimation.

## Glossary

*a*: Hill constant
*b*: Hill constant
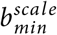: Minimal scale factor of *b* (i.e., the minimal value of *b*_*scale*_)
*b*_*shape*_: Determines the steepness of the relation between *b*_*scale*_ and *q*
*F*_*asymp*_: Oblique asymptote of the eccentric part of the CE force-velocity relationship
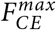: Maximum isometric CE force
*k*_*PEE*_: Scales the PEE stiffness
*k*_*SEE*_: Scales the SEE stiffness
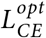: CE length at which CE can produce maximal isometric force
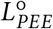: PEE slack length
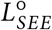: SEE slack length
*r*_*as*_: Slope of the slanted asymptote of the eccentric part of the CE force-velocity relationship
*r*_*slope*_: Ratio between the eccentric and concentric derivatives of 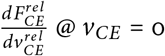
*w*: Determines the width of the CE force-length relationship
*a*_*act*_: Determines the steepness of the relation between *γ* and *q*
*b*_*act*,1_: Determines together with *b*_*act*,1_ and *b*_*act*,3_ the *pCa*^2+^ level at which *q* = 0.5
*b*_*act*,2_: Determines together with *b*_*act*,2_ and *b*_*act*,3_ the *pCa*^2+^ level at which *q* = 0.5
*b*_*act*,3_: Determines together with *b*_*act*,1_ and *b*_*act*,2_ the *pCa*^2+^ level at which *q* = 0.5
*k*_*Ca*_: Relates *γ* to the actual *Ca*^2+^ concentration
*γ*_0_: Minimal value of *γ*
*q*_0_: Minimal value *q*
*τ*_*act*_: Activation time constant of the calcium dynamics
*τ*_*deact*_: Deactivation time constant of the calcium dynamics

## Supplementary material

### S1 Parameter value estimation toolbox

The open-source toolbox automates muscle parameter estimation using data from quick-release, step-ramp, and isometric experiments. Our associated manuscript demonstrates that accuracy improves when correcting for CE shortening during quick-release. However, this requires both quick-release and step-ramp data. Recognising that some users may only have one type of data, we designed the toolbox with flexibility. The toolbox offers the following options:

1. **Estimate CE, SEE and PEE force-length parameters only**
  - Requires *only* quick-release data.
  - Does **not** correct for CE shortening due to the quick-release.
2. **Estimate CE force-velocity parameters only**
  - Requires *only* step-ramp data.
  - Since force-length parameter values are unknown, the following assumptions are made:
    a. CE velocity during ramp equals MTC velocity
    b. CE force equals SEE force
    c. CE length during ramp equals CE optimum length
3. **Estimate force-length and CE force-velocity parameters using the ‘traditional method’**.
  - Requires *both* quick-release and step-ramp data.
  - Does **not** correct for CE shortening due to the quick-release.
4. **Estimate force-length and CE force-velocity parameters using the ‘improved method’** (recommended)
  - Requires *both* quick-release and step-ramp data.
  - **Does** correct for CE shortening due to the quick-release.
5. **Estimate (de-)activation dynamics parameters**
  - Any data type may be used, but isometric contraction data is recommended to minimise the influence of contraction dynamics (see manuscript).
  - Can only be performed after estimating force-length and force-velocity parameters (via option 3 or 4).

The following sections explain how to use each option. Sample scripts are available for all options to further assist with implementation.

#### S1.1 CE, SEE and PEE force-length parameter value estimation

First, extract the relevant data from the quick-release experiments by setting the configurations below:

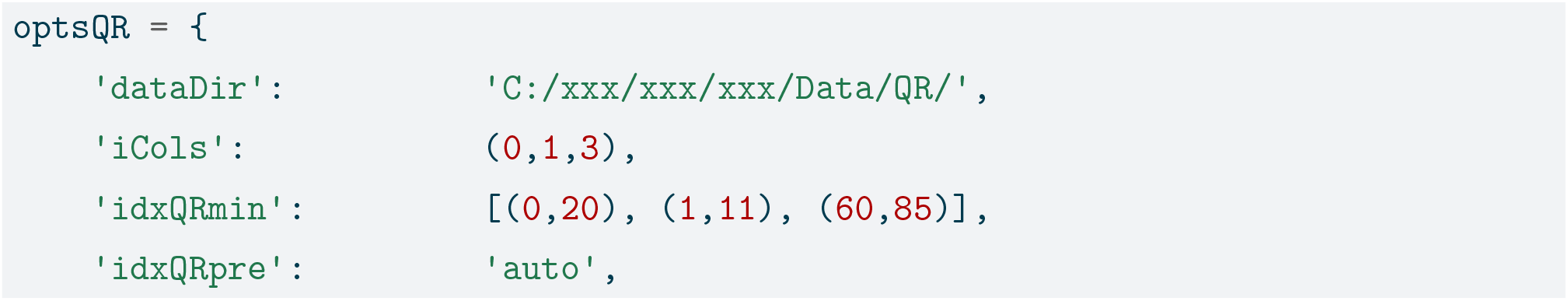
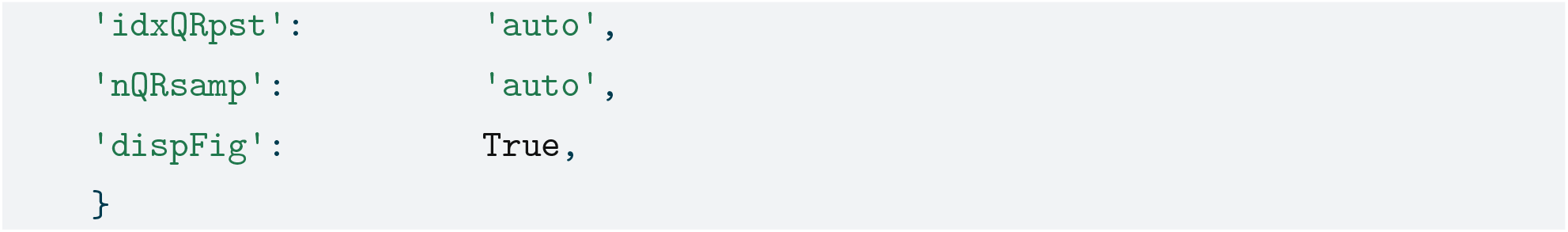

- **dataDir:** Path to your quick-release data files. If omitted, a file selector dialog will appear.
- **iCols:** Tuple indicating the columns representing 1) time; 2 MTC length and 3) SEE force, respectively. If omitted, a column selector dialog will appear.
- **idxQRmin:** List of start and stop indices for the period with minimum SEE force. Provide ranges for each quick-release experiment or use ‘auto’ for automatic detection.
- **idxQRpre:** List of start and stop indices for SEE force before the quick-release. Provide ranges for each quick-release experiment or use ‘auto’ for automatic detection.
- **idxQRpst:** List of start and stop indices for SEE force after the quick-release. Provide ranges for each quick-release experiment or use ‘auto’ for automatic detection.
- **nQRsamp:** Number of samples to average before and after the quick-release. Default is 5 if ‘auto’ or unspecified.
- **dispFig:** When True, plots will be displayed for manual verification and adjustment of selected data segments (i.e., idxQRmin, idxQRpre and idxQRpst).

Second, call the function ‘loaddata.qr’ to select the quick-release data:

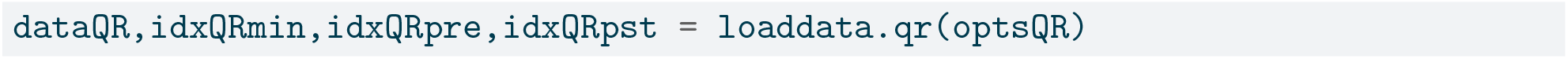

- **dataQR:** List of dictionaries containing the extracted quick-release data.
- **idxQRmin, idxQRpre and idxQRpst:** Returned index ranges for your reference or future use.

Third, obtain the ‘default’ muscle parameter (such as width of the CE force-length relationship etc.):

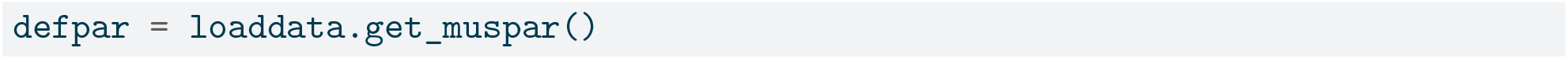

- **defpar:** Dictionary containing the ‘default’ muscle parameter values.

Fourth, estimate the CE, PEE and SEE and force-length parameter values by passing your data and initial parameters to ‘estimate.fl’:

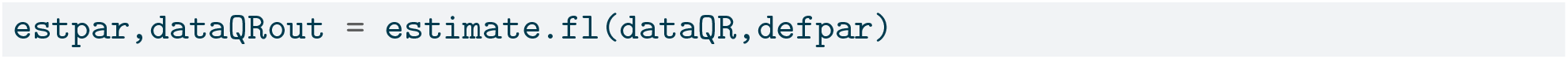

- **estpar:** Dictionary containing estimated muscle parameter values.
- **dataQRout:** List of dictionaries for each quick-release trial, with keys:
  - *lseeQRpre:* Estimated SEE length before quick-release
  - *lseeQRpst:* Estimated SEE length after quick-release
  - *fseeQRpre:* SEE force before quick-release
  - *fseeQRpst:* SEE force after quick-release
  - *lpeeQR:* (Estimated) PEE length of the data
  - *fpeeQR:* PEE force of the data
  - *lmtcQRpre:* MTC length before quick-release

#### S1.2 CE force-velocity parameter value estimation

First, extract the relevant data from the step-ramp experiments by setting the configurations below:

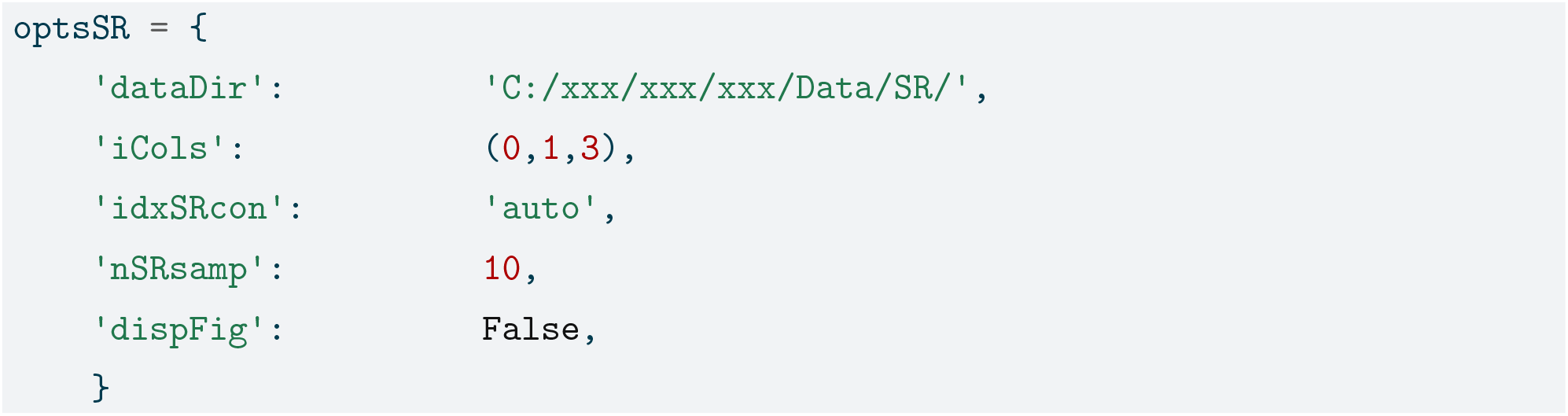

- **dataDir:** Path to your step-ramp data files. If omitted, a file selector dialog will appear.
- **iCols:** Tuple indicating the columns representing 1) time; 2 MTC length and 3) SEE force, respectively. If omitted, a column selector dialog will appear.
- **idxSRcon:** List of start and stop indices where SEE force is most ‘constant’. If set to ‘auto’, indices are automatically detected.
- **nSRsamp:** Number of data points to average during the constant force phase. Default is 10 if ‘auto’ or unspecified.
- **dispFig:** When True, plots will be displayed for manual verification and adjustment of selected data segments (i.e., idxSRcon).

Second, call the function ‘loaddata.sr’ to select the step-ramp data:

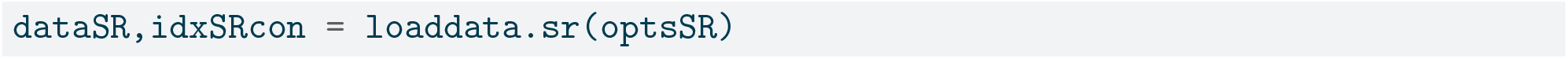

- **dataSR:** List of dictionaries containing the extracted step-ramp data.
- **idxSRcon:** Returned index ranges for your reference or future use.

Third, obtain the ‘default’ muscle parameter (such as width of the CE force-length relationship etc.):

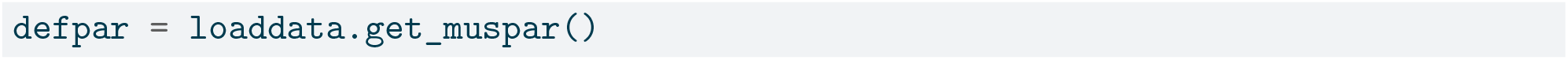

- **defpar:** Dictionary containing the ‘default’ muscle parameter values.

Fourth, estimate the CE force-velocity parameter values by passing your data and initial parameters to ‘estimate.fv’:

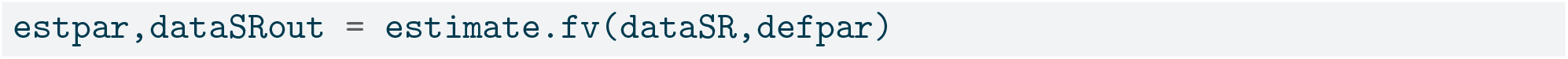

- **estpar:** Dictionary containing the estimated muscle parameters.
  - If only CE force-velocity parameters are estimated (option 2), *a, b*, and 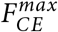 are estimated on based step-ramp data.
  - If combined with estimation of force-length parameters (option 3 & 4), *a, b* are estimated on based step-ramp data (while 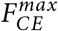 is estimated based on quick-release data).
- **dataQRout:** List of dictionaries for each quick-release trial, with keys:
  - *vceSR:* Estimated CE shortening velocity.
  - *fceSR:* Estimated CE force.
  - *lcerelSR:* Relative CE length.

#### S1.3 Traditional method

Running the traditional method requires first estimating the CE, SEE and PEE force-length parameter values and then CE force-velocity parameter values. All inputs and outputs of the function are described above.

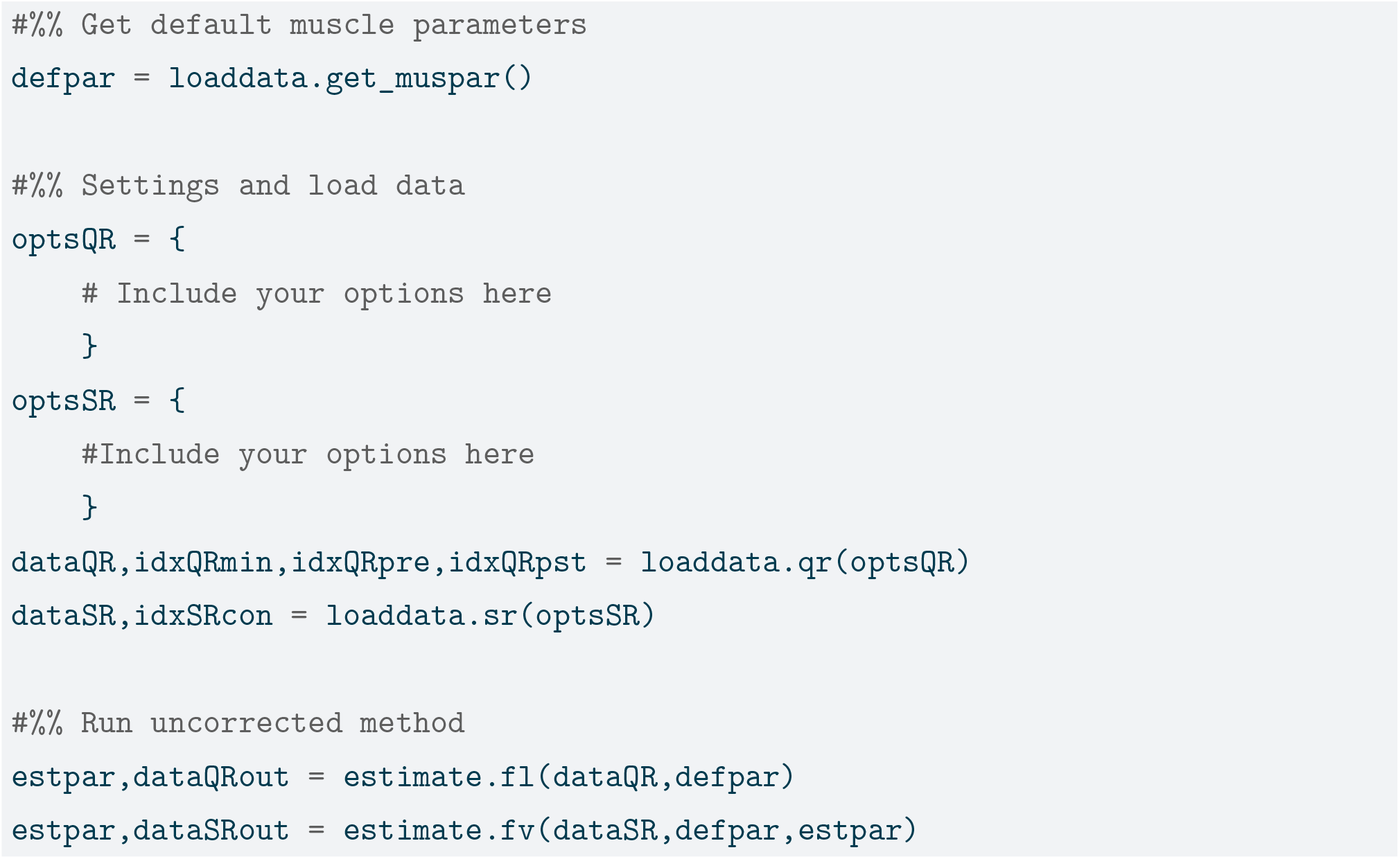

#### S1.4 Improved method

To run the improved method, call a separate function ‘estimate.im’. This function iteratively call ‘estimate.fl’ and ‘estimate.fv’ until the change in all estimated parameter values are below 0.1%. All inputs and outputs of the function are described above.

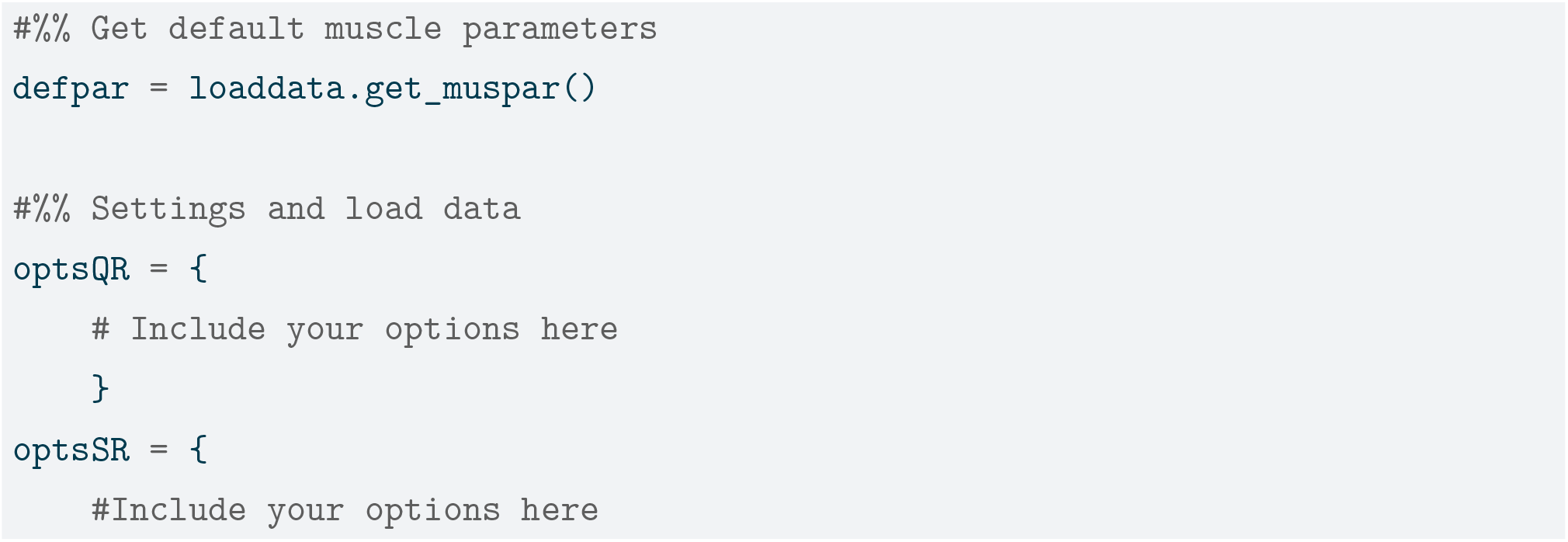
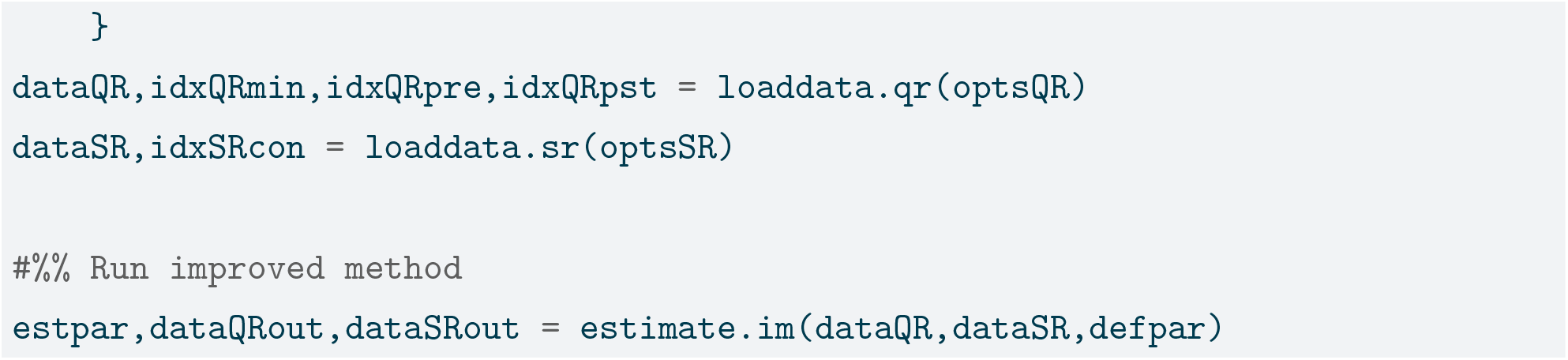

#### S1.5 Excitation dynamics parameter value estimation

To estimate the time constants of the calcium dynamics, one has to first estimate the parameter values of the CE, PEE and SEE force-length relationships as well as the parameters of the CE force-velocity relationship. After this the time constant of the calcium dynamics can be estimated. First, extract the relevant data (from the quick-release experiments) by setting the configurations below:

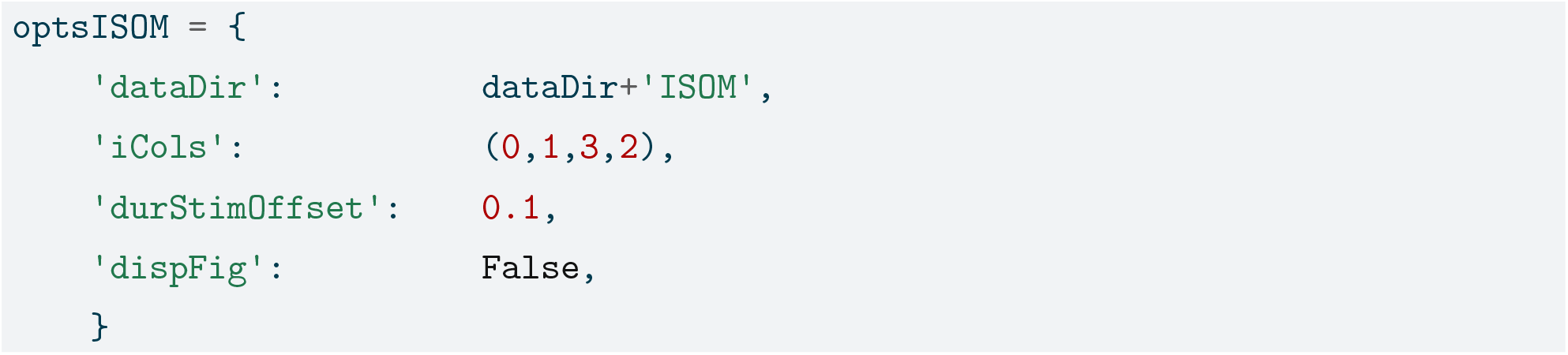

- **dataDir:** Path to your (isometric) data files. If omitted, a file selector dialog will appear.
- **iCols:** Tuple indicating the columns representing 1) time; 2 MTC length and 3) SEE force, respectively. If omitted, a column selector dialog will appear.
- **durStimOffset:** Time (in seconds) to include after stimulation offset. If not specified, the default is 0.1 s.
- **dispFig:** When True, plots will be displayed for manual verification of selected data segments (i.e., idxSRcon).

Second, call the function ‘SelISOMdata’ to select the (isometric) data:

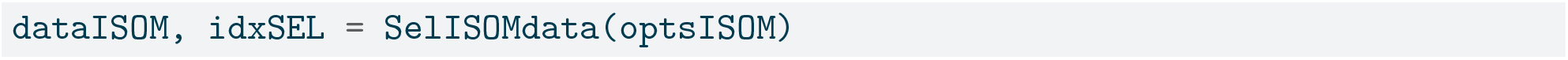

- **dataISOM:** List of dictionaries containing the extracted (isometric) data.
- **idxSEL:** List of start and stop indices of selected interval of the data.

Third, estimate the excitation dynamics parameter values by passing your data and initial parameters to ‘getACTparms’:

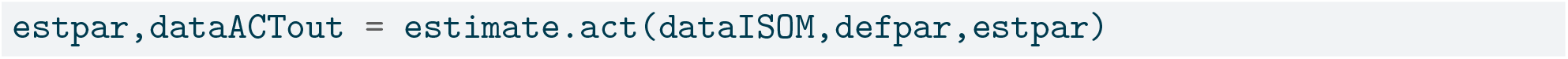

- **estpar:** Dictionary containing the estimated muscle parameters.
  - **dataACTout:** List of dictionaries for each (isometric) experment, with keys::
  - *time:* Time-axis.
  - *lmtc:* MTC length.
  - *fseeData:* Experimental SEE force.
  - *fseeMdl:* Model-predicted SEE force.
  - *tStim:* List containing stimulation onset and offset times.

### S2 Supplemental Figures

**Figure S1:**
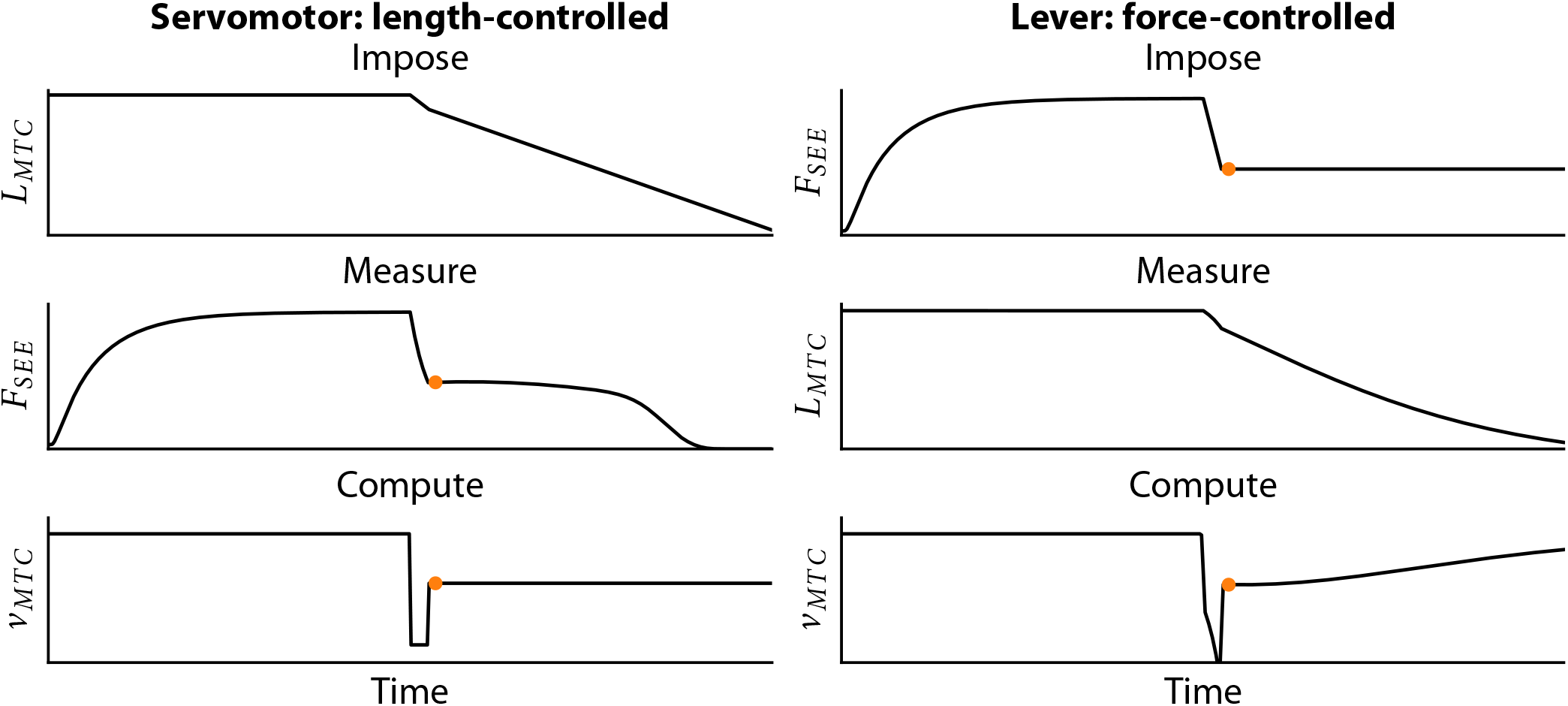
Comparison between a length-controlled step-ramp experiment and a force-controlled quick-release experiment. In length-controlled step-ramp experiment (left), MTC length over time is imposed using a servomotor, while SEE force is measured. MTC velocity is computed at the time instance at which SEE force is near constant. In the force-controlled quick-release experiment (right), SEE force over time is imposed via a lever, while MTC length is measured. MTC velocity is computed at the time instance where MTC velocity is maximal after the change in SEE force. Both methods yield one datapoint (depicted with the orange dot) of the force-velocity relationship.

**Figure S2:**
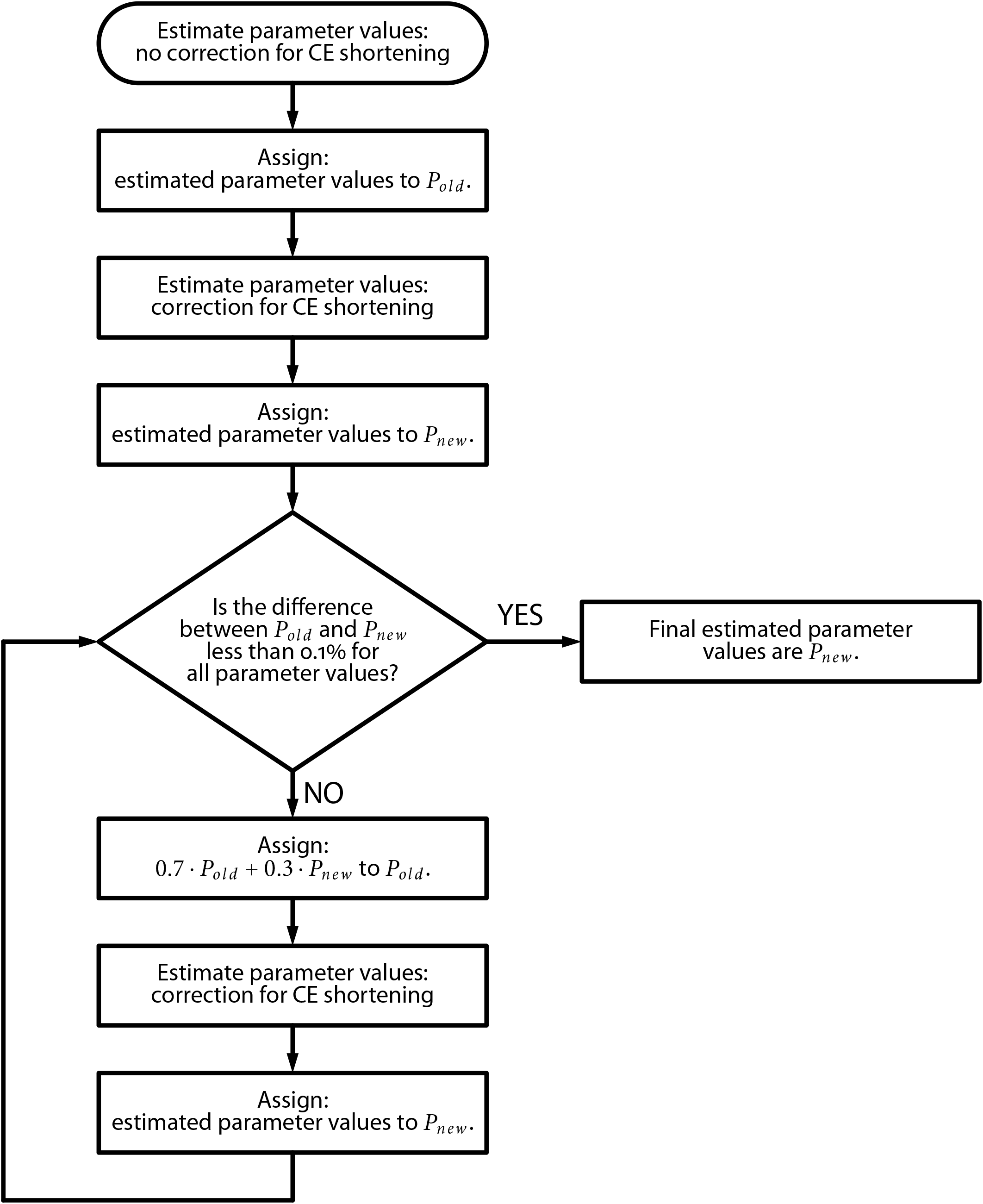
Flowchart of the improved method. Using the improved method, parameter values were estimated until the change in all parameter values was less than 0.1%.

**Figure S3:**
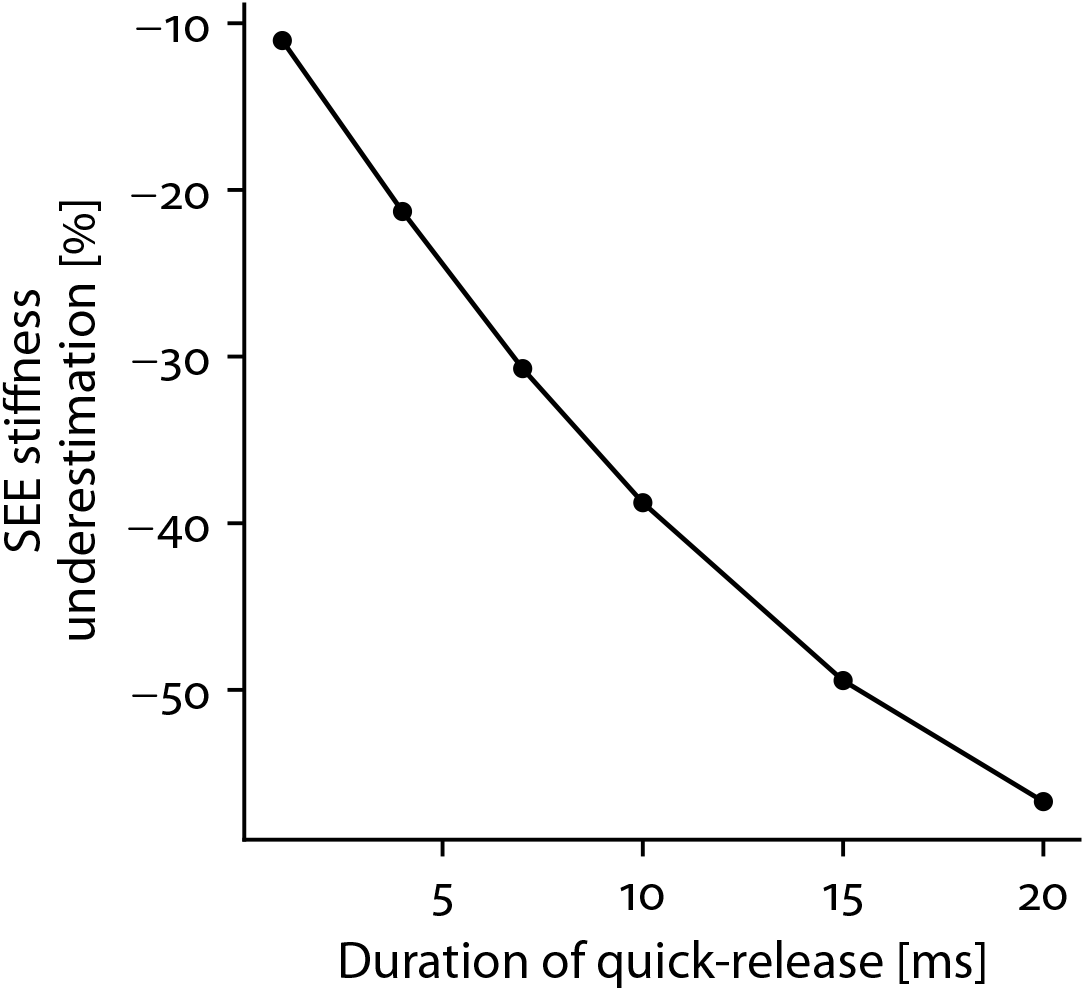
Relationship between quick-release duration and SEE stiffness understimation. When the duration of the quick-release increases, the underestimation of SEE stiffness increases.

### S3 Supplemental Tables

**Table S1:** MTC parameter values obtained from literature to simulate data.

| Parameter | Unit | GM1 | GM2 | GM3 | Reference |
| --- | --- | --- | --- | --- | --- |
| *Contraction dynamics* |  |  |  |  |  |
| $a$ | N | 2.68 | 1.80 | 2.58 | van Zandwijk et al. (1996) |
| $b$ | mm | 41.6 | 24.8 | 41.8 | van Zandwijk et al. (1996) |
| $b_{scale}^{min}$ | - | 0.100 | | | van Soest and Bobbert (1993) |
| $b_{shape}$ | - | 22.0 | | | van Soest and Bobbert (1993) |
| $F_{asympt}$ | - | 1.50 | | | Rijkelhuizen et al. (2003) |
| $F_{CE}^{max}$ | N | 13.4 | 13.8 | 12.3 | van Zandwijk et al. (1996) |
| $k_{PEE}$ | N/mm<sup>2</sup> | 213 | 165 | 511 | van Zandwijk et al. (1996) |
| $k_{SEE}$ | N/mm<sup>2</sup> | 4220 | 3640 | 3470 | van Zandwijk et al. (1996) |
| $L_{CE}^{opt}$ | mm | 13.2 | 12.3 | 11.2 | van Zandwijk et al. (1996) |
| $L_{PEE}^o$ | mm | 13.9 | 13.2 | 13.4 | van Zandwijk et al. (1996) |
| $L_{SEE}^o$ | mm | 28.3 | 30.3 | 26.5 | van Zandwijk et al. (1996) |
| $r_{as}$ | - | $3.71 \cdot 10^{-3}$ | $5.45 \cdot 10^{-3}$ | $3.16 \cdot 10^{-3}$ | Arbitrary small value |
| *Excitation dynamics* |  |  |  |  |  |
| $r_{slope}$ | mm | 2.00 | | | Katz (1939) |
| $w$ | - | 0.50 | | | Burkholder and Lieber (2001) |
| $a_{act}$ | - | -7.37 | | | Bortolotto et al. (2000) |
| $b_{act,1}$ | - | 5.17 | | | Bortolotto et al. (2000) |
| $b_{act,2}$ | - | 0.596 | | | Stephenson and Williams (1982) |
| $b_{act,3}$ | - | 0.00 | | | Stephenson and Williams (1982) |
| $kCa$ | mol/L | $8.00 \cdot 10^{-6}$ | | | Kistemaker et al. (2005) |
| $\gamma_o$ | - | $1.00 \cdot 10^{-5}$ | | | Arbitrary small value |
| $q_o$ | - | $5.00 \cdot 10^{-3}$ | | | Hatze (1981) |
| $\tau_{act}$ | ms | 27.0 | | | van Zandwijk et al. (1996) |
| $\tau_{deact}$ | ms | 27.0 | | | van Zandwijk et al. (1996) |

**Table S2:** Estimated parameter values of the experimentally measured *in situ* data using both the traditional as well as the improved method.

|  |  | *Traditional method* |  |  | *Improved method* |  |  |
| --- | --- | --- | --- | --- | --- | --- | --- |
| **Parameter** | **Unit** | **Rat 1** | **Rat 2** | **Rat 3** | **Rat 1** | **Rat 2** | **Rat 3** |
| $a$ | N | 8.90 | 11.3 | 10.5 | 8.82 | 10.0 | 11.0 |
| $b$ | mm | 76.7 | 108 | 84.6 | 81.9 | 105 | 90.6 |
| $F_{CE}^{max}$ | N | 15.6 | 14.1 | 17.1 | 15.5 | 14.1 | 17.0 |
| $k_{PEE}$ | N/mm<sup>2</sup> | 9.83 | 7.25 | 21.9 | 28.7 | 24.1 | 34.8 |
| $k_{SEE}$ | N/mm<sup>2</sup> | 608 | 688 | 654 | 952 | 1260 | 1050 |
| $L_{CE}^{opt}$ | mm | 12.4 | 12.9 | 12.3 | 13.7 | 14.7 | 13.3 |
| $L_{PEE}^o$ | mm | 12.4 | 12.4 | 13.3 | 15.1 | 15.6 | 14.5 |
| $L_{SEE}^o$ | mm | 29.0 | 29.7 | 25.1 | 28.8 | 29.3 | 25.2 |
| $\tau_{act}$ | ms | 43.0 | 32.4 | 23.6 | 55.2 | 57.7 | 41.6 |
| $\tau_{deact}$ | ms | 23.8 | 20.9 | 19.7 | 27.1 | 25.3 | 22.4 |

**Table S3:** Root mean squared differences between experimentally measured SEE force histories and those predicted by Hill-type MTC model after estimating all contraction and excitation dynamics parameter values.

|  | *Traditional method* |  |  |  | *Improved method* |  |  |  |
| --- | --- | --- | --- | --- | --- | --- | --- | --- |
|  | **QR** | **SR** | **ISOM** | **SSC** | **QR** | **SR** | **ISOM** | **SSC** |
| Rat 1 | 5.6 ± 3.4 | 5.6 ± 0.8 | 6.8 ± 3.2 | 6.2 ± 3.0 | 3.3 ± 1.5 | 2.8 ± 0.9 | 5.4 ± 3.0 | 5.4 ± 3.1 |
| Rat 2 | 4.2 ± 1.5 | 5.0 ± 0.6 | 4.3 ± 1.6 | 4.6 ± 2.2 | 2.7 ± 1.6 | 2.1 ± 0.5 | 4.2 ± 2.6 | 4.2 ± 2.0 |
| Rat 3 | 6.4 ± 2.3 | 6.8 ± 1.0 | 6.1 ± 2.1 | 5.5 ± 2.7 | 4.2 ± 1.8 | 2.6 ± 0.5 | 4.9 ± 2.3 | 5.0 ± 2.7 |
| Avg ± Std | 5.4 ± 2.6 | 5.8 ± 1.1 | 5.7 ± 2.7 | 5.4 ± 2.7 | 3.4 ± 1.8 | 2.5 ± 0.7 | 4.8 ± 2.7 | 4.9 ± 2.7 |

Root mean squared differences are expressed as a percentage of the maximal isometric CE force ( $F_{CE}^{max}$ ). Root mean squared differences were computed over the interval in which CE stimulation was maximal for the quick-release and step-ramp experiments and was computed over the interval from CE stimulation onset to 0.1 s after CE stimulation ceased for the isometric experiments and the stretch-shortening cycles.

